## Supplementary information for "Engineering of a fluorescent chemogenetic reporter with tunable color for advanced live-cell imaging"

**This PDF file includes:**

Legends for Movies S1 to S7
Texts S1 to S2
Figures S1 to S19
Tables S1 to S13
SI References

**Other supplementary materials for this manuscript include the following:**

Movies S1 to S7

**Legends for Movies**

**Movie S1. Labeling efficiency of pFAST and FAST with HBR-3,5DOM in dissected neural tube of chicken embryo.** Plasmids encoding H2B-pFAST and H2B-FAST were each electroporated in one of each side of the neural tube in ovo at embryonic day 2 (E2, HH stage 13-14). An EGFP reporter fused to H2B was co-injected with each construct to monitor electroporation efficiency. 24 h later, embryos with homogeneous bilateral EGFP expression in the neural tube were dissected, and imaged upon successive additions of 0.1, 1 and 10 μM HBR-3,5DOM using a spinning-disk confocal microscope (see **Table S13** for imaging settings).

**Movie S2. Two-color dynamic cellular processes in dissected neural tube of chicken embryo.** Plasmids encoding mito-pFAST (fused to a mitochondrial localization signal) and mb-mIRFP670 (fused to a membrane localization signal) were electroporated in the neural tube in ovo, at embryonic day 2 (E2, HH stage 13-14). 24 h later, embryos were dissected, and imaged in presence of 1 μM HBR-3,5DOM using a spinning-disk confocal microscope (see **Table S13** for imaging settings). The time-lapse shows cell division. Scale bar, 10 μm.

**Movie S3. Two-color imaging in dissected neural tube of chicken embryo.** Plasmids encoding H2B-pFAST (fused to histone H2B) and pact-mKO (targeting the PACT domain of pericentrin) were electroporated in the neural tube in ovo, at embryonic day 2 (E2, HH stage 13-14). 24 h later, embryos were dissected, and imaged in presence of 5 μM HBP-3,5DOM using a spinning-disk confocal microscope (see **Table S13** for imaging settings). The time-lapse shows cell division. Scale bar, 10 μm.

**Movie S4. Three-color imaging in dissected neural tube of chicken embryo.** Plasmids encoding H2B-pFAST (fused to histone H2B), pact-mKO (targeting the PACT domain of pericentrin) and mb-miRFP670 (fused to a membrane localization signal) were electroporated in the neural tube in ovo at embryonic day 2 (E2, HH stage 13-14). 24 h later, embryos were dissected, and imaged in presence of 1 μM HMBR using a spinning-disk confocal microscope. The time-lapse shows cell division (see **Table S13** for imaging settings). Scale bar, 10 μm.

**Movie S5. Reversible labeling of pFAST in live mammalian cells by chromophore replacement**. Time-lapse imaging of HeLa cells expressing cytoplasmic pFAST initially labeled with 1 μM HBR-3,5DM, 10 μM HBP-3,5DOM, 1 μM HMBR and 5 μM HBP-3,5DM were switched off upon addition of 10 μM HBIR-3M dark-competitor (the concentration of fluorogenic chromophore was kept constant during the overall experiment). The transmitted channels allowed to visualize the cells (see **Table S13** for imaging settings). Scale bars, 30 μm.

**Movie S6. Reversible labeling of pFAST in dissected neural tube of chicken embryo.** Plasmid encoding H2B-pFAST was electroporated in the neural tube in ovo at embryonic day 2 (E2, HH stage 13-14). An mRFP reporter fused to H2B was co-injected to monitor electroporation efficiency. 24 h later, embryos with homogeneous bilateral mRFP expression in the neural tube were dissected, and initially labeled with 1 μM HMBR for 40 minutes. Time-lapse imaging allowed to monitor the fluorescence evolution after washing the embryos with PBS and addition of fresh medium supplemented with 0 μM ("wash"), 1 μM and 10 μM of the dark competitor HBIR-3M prior to imaging (see **Table S13** for imaging settings).

**Movie S7. Rapid dynamics of microtubules and membranes in live cells observed with high temporal and spatial resolution by Airyscan confocal microscopy.** Time-lapse imaging of HeLa cells expressing lyn11-pFAST (fused to membrane localization signal) (left) and MAP4-pFAST (microtubule associated protein) (right) and labeled with 5 μM HBR-3,5DOM allowed to visualize the rapid dynamics of membrane and microtubules with high temporal and spatial resolution by Airyscan confocal microscopy (see **Table S13** for imaging settings). Scale bars, 10 μm.

**Text S1: Full description of the directed protein evolution experiments leading to oFAST, tFAST and pFAST**

This supplementary text complements the main text, and details the full directed evolution experiments done during this study.

We used a combinatorial library of $10^6$ variants of FAST generated by random mutagenesis and displayed on yeast cells (library A). Six different screenings in presence of either HBO-3M, HBO-3,5DM, HBT-3M, HBT-3,5DM, HBP-3M or HBP-3,5DM were performed by iterative rounds of fluorescence activated cell sorting (FACS) decreasing progressively chromophore concentrations through rounds (typically from 10 μM to 2.5 μM) in order to identify variants forming tighter and brighter assemblies (**Fig. S2**, **S3**). The screenings with HBO-3M, HBO-3,5DM and HBP-3,5DM showed an increase in cell population fluorescence indicating the selection of improved variants (**Tables S5-S7**). For each positive screening, we systematically isolated and sequenced twenty-four clones after the fifth, sixth and seventh round of FACS, and then further analyzed the performances of single clones by analytical flow cytometry. Five to ten individual clones with the highest fluorescence performances relative to FAST were expressed in bacteria and purified for in vitro characterization. The variants isolated from the three screenings formed tighter and brighter assemblies with the chromophore used for their selection (**Fig. 2b,d**, **Fig. S4a**, **Tables S5-7**), in agreement with a successful molecular evolution.

By screening variants that combined mutations beneficial for the binding of HBO-3M (**Table S5**) and HBO-3,5DM (**Table S6**), we identified oFAST, an improved variant with the mutations Q41L, D71N, V83I, M109L and S117R, showing improved properties with both HBO-3M and HBO-3,5DM. oFAST binds HBO-3,5DM with a $K_D$ of 3.0 μM, forming a blue fluorescent complex with 411/482 nm abs/em peaks and a fluorescent quantum yield $\phi$ = 30%, and it binds HBO-3M more tightly with a $K_D$ of 0.73 μM, forming a blue fluorescent complex with 394/470 nm abs/em peaks and a fluorescent quantum yield $\phi$ = 11% (**Table S2**).

The third positive selection allowed us to identify several variants able to bind HBP-3,5DM with 3 to 7 fold higher affinity, and giving complexes displaying higher fluorescence quantum yield. Beneficial mutations isolated from the three brighter variants were combined, leading eventually to three new variants binding HBP-3,5DM about 10-fold tighter than FAST and leading to higher fluorescent quantum yields (from $\phi$ = 22 to 26 %) (**Table S7**). The five more promising variants were further characterized with the other members of the HBP series and with the chromophores of the HBT series. We discovered that the five variants were also able to form tighter and brighter assemblies with HBT-3,5DM (**Table S8**), HBT-3,5DOM, HBP-3M and HBP-3,5DOM (**Table S9**).

We thus used these five variant genes to construct a new library using DNA shuffling, which allows random DNA fragments recombination to access positive combinations. The five variant genes were randomly recombined to generate a library of $5.5 \times 10^7$ variants (library B), which was displayed on yeast cells. This new library was screened in presence of HBT-3,5DM, HBP-3,5DM and HBP-3,5DOM by iterative rounds of FACS as described above. The screening with HBP-3,5DOM did not allow us to identify variants with significantly better properties (**Fig. S4b**, **Table S9**). However, the screening with HBT-3,5DM allowed us to identify variants able to form brighter complexes (**Fig. 2c**, **Table S8**). The variant possessing the mutations G25R, Q41K, S72T, A84S, M95A, M109L and S117R relative to FAST, which we ultimately named tFAST, showed the most advantageous properties: it was shown to form a tight cyan fluorescent assembly with HBT-3,5DM ($K_D$ = 0.33 μM, 435/497 nm abs/em peaks, $\phi$ = 11%) (**Table S3**). On the other hand, the screening of the library B with HBP-3,5DM permitted us to identify an improved variant bearing the mutations K17N, G21E, G25R, A30V, Q41L, S72T, V83A, M95T, S117R, which was further refined by introduction of the mutation M109L found in several other improved variants. The resulting variant, named pFAST, binds HBP-3,5DM tightly with a $K_D$ of 0.15 μM forming a bright green fluorescent assembly ($\phi$ = 27%, 465/520 nm abs/em peaks) (**Table S4**, **Table S7**).

**Text S2: Theoretical model for the determination of the thermodynamic dissociation constants of the dark chromophores**

The thermodynamic dissociation constants ($K_D$) of the dark HBIR-3M and HBIR-3,5DM chromophores were determined by determining the apparent dissociation constant of HBP-3,5DM in presence of various concentrations of dark competitors. Considering that the fluorogen HBP-3,5DM and the protein interact to provide a fluorescent complex whereas the dark chromophores HBIR-3M or HBIR-3,5DM and the protein interact to form a non-fluorescent complex, we adopted the following three-state model

$$\begin{array}{ccc} F_1 + & R & \leftrightarrows F_1R \\ & + & \\ & F_2 & \\ & \uparrow\downarrow & \\ & F_2R & \end{array}$$

where $F_1$ denotes the fluorogen, $F_2$ denotes the dark chromophore, R denotes the protein and $F_1R$ and $F_2R$ denote the two possible complexes, characterized by the dissociation constants

$$K_{D,1} = \frac{[F_1]\ [R]}{[F_1R]}$$

and

$$K_{D,2} = \frac{[F_2]\ [R]}{[F_2R]}$$

where [X] is the concentration of the species X at equilibrium. At equilibrium, the fraction of protein R binding $F_1$ is given by

$$B = \frac{[F_1R]}{[R] + [F_1R] + [F_2R]} = \frac{\frac{[F_1]\ [R]}{K_{D,1}}}{[R] + \frac{[F_1]\ [R]}{K_{D,1}} + \frac{[F_2]\ [R]}{K_{D,2}}} = \frac{[F_1]}{[F_1] + K_{D,1}\left(1 + \frac{[F_2]}{K_{D,2}}\right)}$$

$$= \frac{[F_1]}{[F_1] + K_{D,app}}$$

with

$$K_{D,app} = K_{D,1}\left(1 + \frac{[F_2]}{K_{D,2}}\right)$$

By choosing $[F_1]_0$ and $[F_2]_0 >> [R]_0$, one can rewrite this equation assuming that $[F_1] \approx [F_1]_0$ and $[F_2] \approx [F_2]_0$. Knowing $K_{D,1}$, determination of the apparent thermodynamic dissociation constant $K_{D,app}$ at various $[F_2]_0$ enabled to determine $K_{D,2}$.

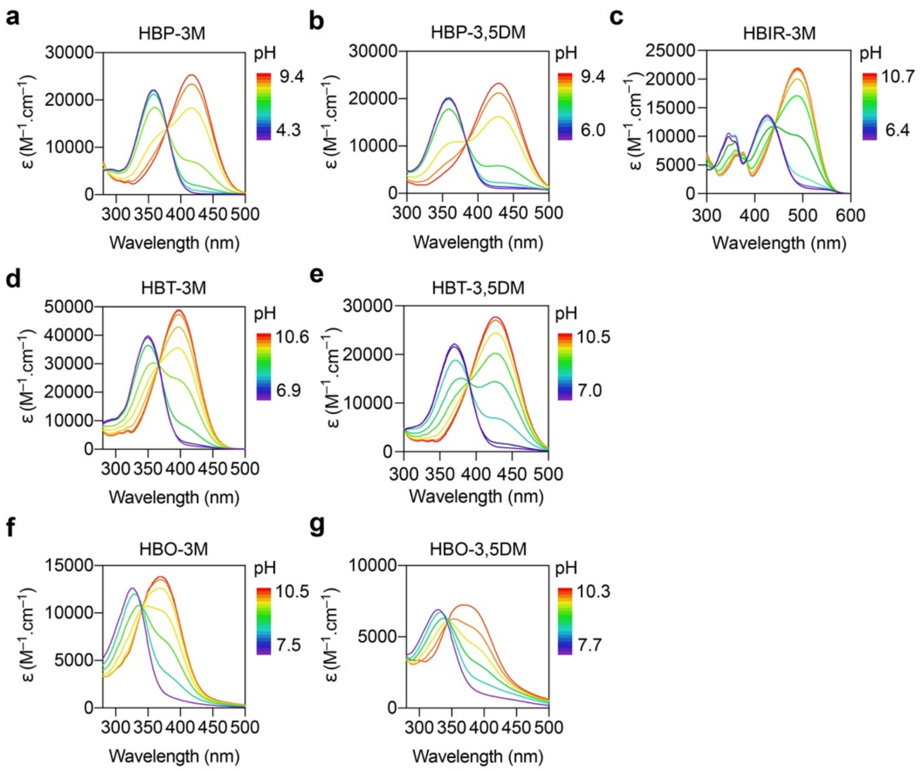


**h**

| Fluorogens | p$K_A$ |
|---|---|
| HBP-3M | 8.15 |
| HBP-3,5DM | 8.25 |
| HBT-3M | 8.70 |
| HBT-3,5DM | 8.80 |
| HBO-3M | 8.90 |
| HBO-3,5DM | 9.05 |
| HBIR-3M | 8.29 |

**Fig. S1. Absorption spectra of fluorogenic chromophore at various pH.** Absorption spectra of **(a)** HBP-3M, **(b)** HBP-3,5DM, **(c)** HBIR-3M, **(d)** HBT-3M, **(e)** HBT-3,5DM, **(f)** HBO-3M and **(g)** HBO-3,5DM in solution in function of pH. The spectra were recorded in 0.01 M Britton-Robinson buffer[1] (0.1 M ionic strength) at 25°C. **(h)** Extracted p$K_A$ values.

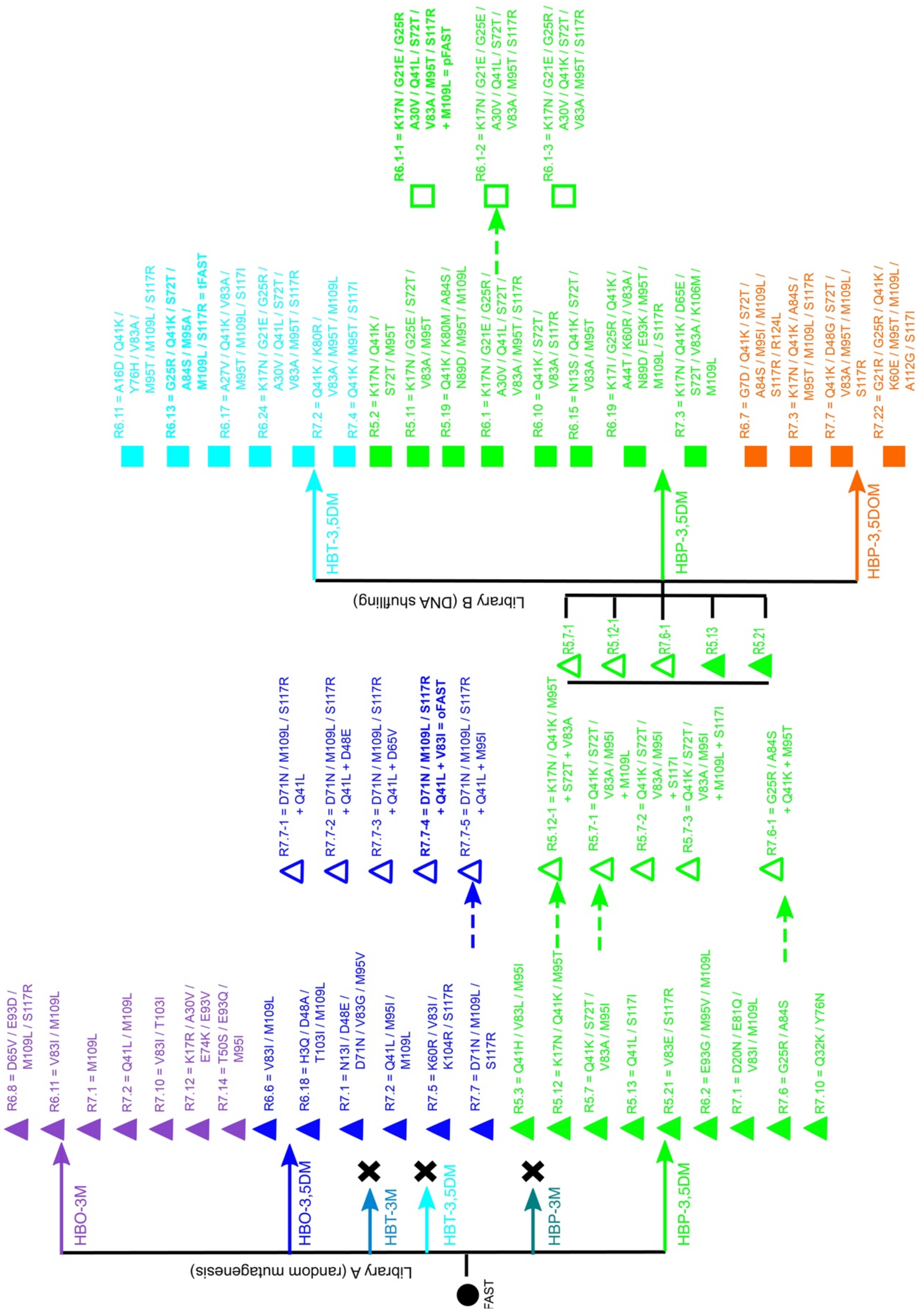


**Fig. S2. Evolutionary tree of the selections with HBO-3M (in purple), HBO-3,5DM (in blue), HBT-3,5DM (in cyan), HBP-3,5DM (in green) and HBP-3,5DOM (in orange).** Selected clones were generated by directed evolution from initial libraries of mutants (solid arrows). The first library was constructed by random mutagenesis from the original FAST sequence (library A) and the second library was designed by DNA shuffling from five mutants, initially selected and rationally designed from the selection for HBP-3,5DM binders (library B). Additional mutations were introduced by rational design (dotted arrows). The crosses indicate unsuccessful selections. The mutations relative to FAST are indicated for each clone.

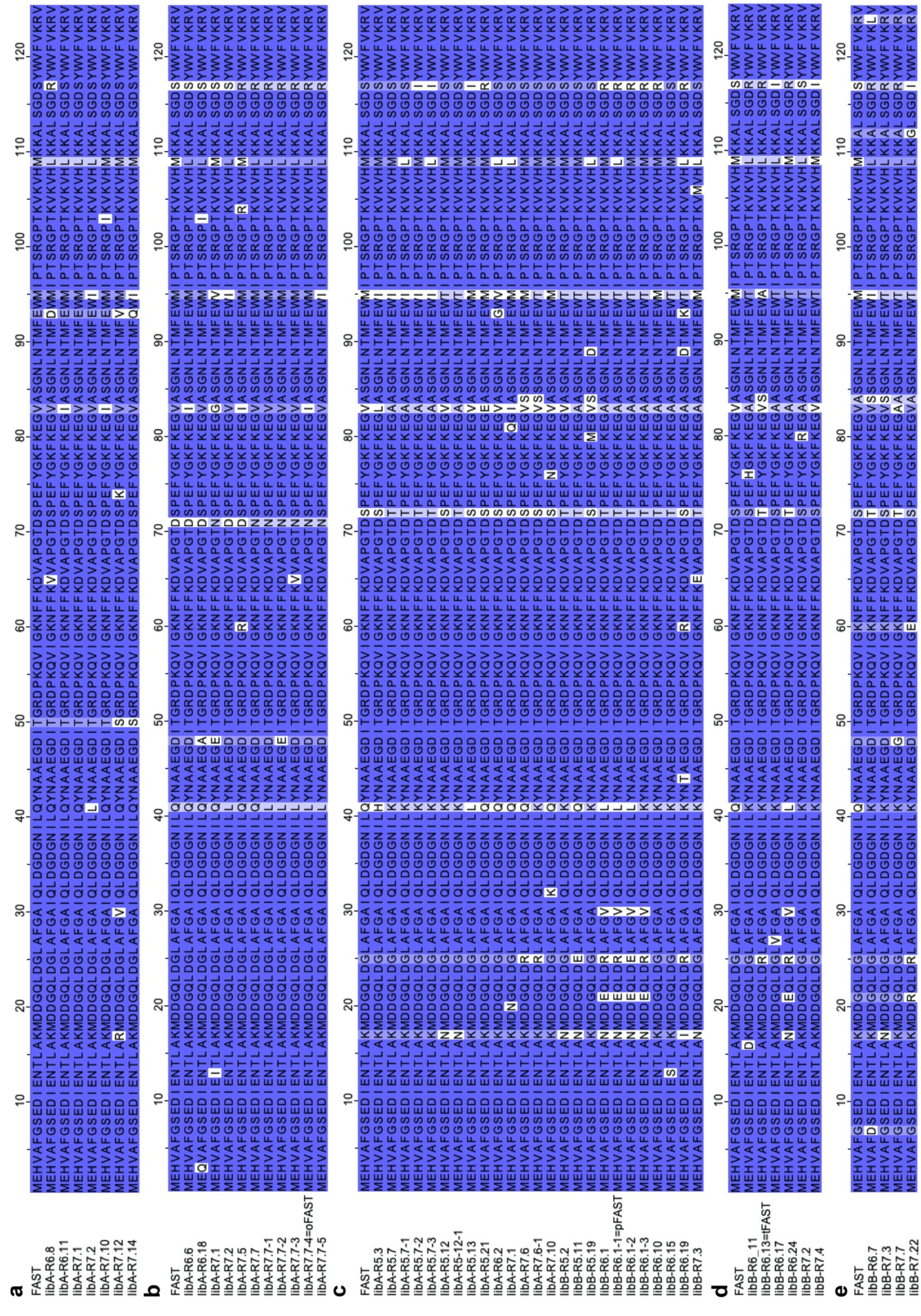

**Fig. S3.** Sequence alignment of FAST and the clones identified from the selections with **(a)** HBO-3M, **(b)** HBO-3,5DM, **(c)** HBP-3,5DM, **(d)** HBT-3,5DM and **(e)** HBP-3,5DOM.

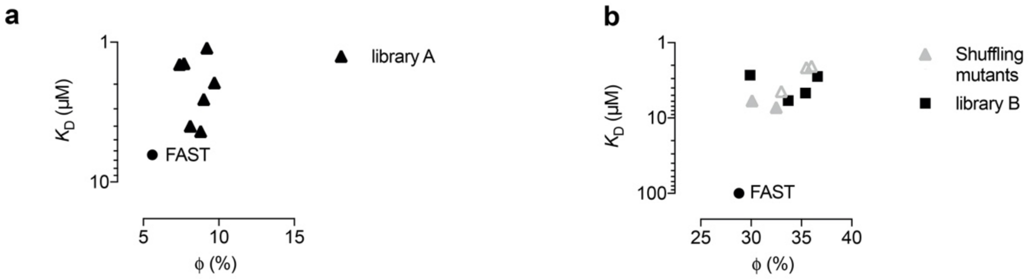


**Fig. S4. Characteristics of the selected clones.** Thermodynamic dissociation constants ($K_D$) and fluorescent quantum yields ($\phi$) of the clones isolated from the selections with **(a)** HBO-3M and **(b)** HBP-3,5DOM. Values are also given for FAST for comparison. **(a)** Mutants of HBO-3M selection were isolated from the library A (black triangles). **(b)** Mutants of HBP-3,5DOM were isolated from the library B (black squares) and compared to the shuffling mutants used to design the library B (gray triangles).

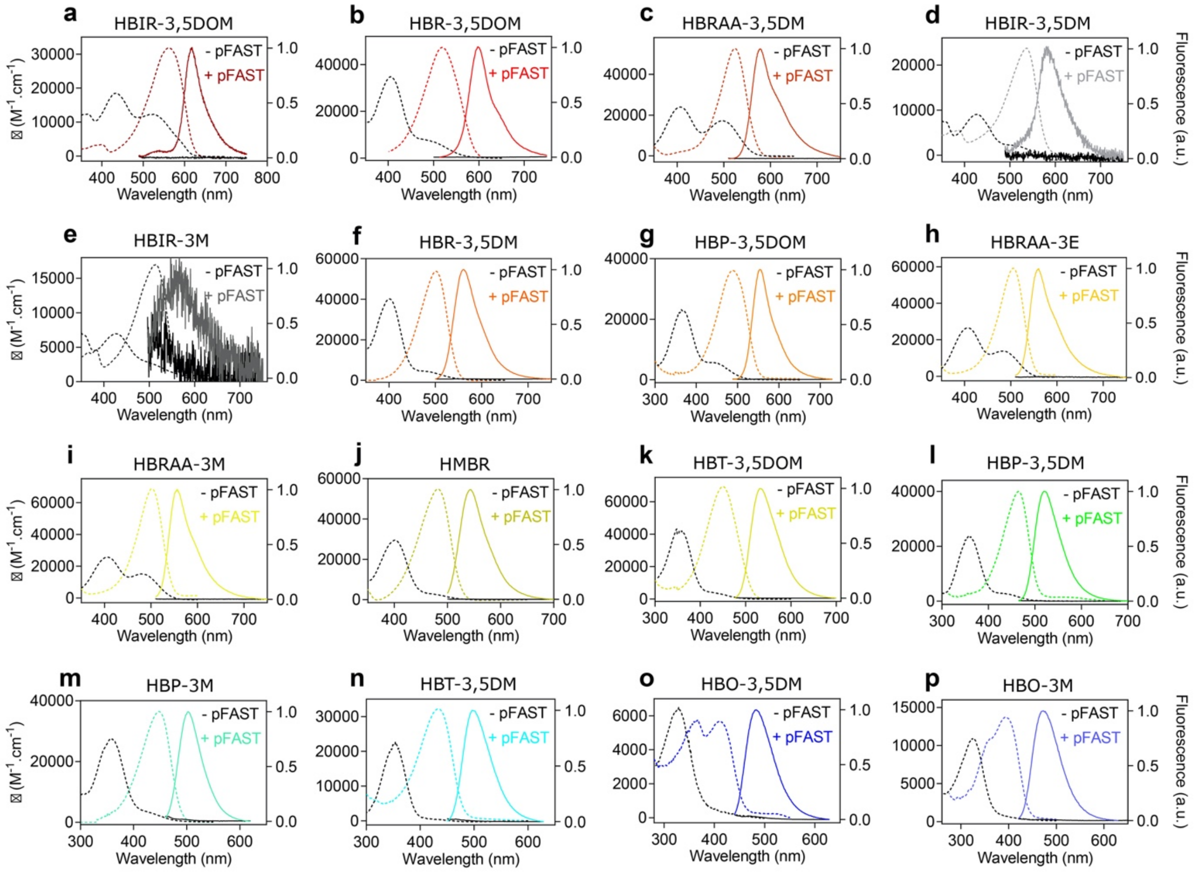


**Fig. S5. Absorption and emission properties.** Absorption (dashed lines) and fluorescence (solid lines) spectra of the fluorogenic chromophores when free in solution (dark line) and bound to pFAST (colored lines) for **(a)** HBIR-3,5DOM, **(b)** HBR-3,5DOM, **(c)** HBRAA-3,5DM, **(d)** HBIR-3,5DM, **(e)** HBIR-3M, **(f)** HBR-3,5DM, **(g)** HBP-3,5DOM, **(h)** HBRAA-3E, **(i)** HBRAA-3M, **(j)** HMBR, **(k)** HBT-3,5DOM, **(l)** HBP-3,5DM, **(m)** HBP-3M, **(n)** HBT-3,5DM, **(o)** HBO-3,5DM and **(p)** HBO-3M. Spectra were recorded using chromophore concentrations ranging from 3 – 15 µM and pFAST at 40 µM in pH 7.4 phosphate buffer saline at 25°C.

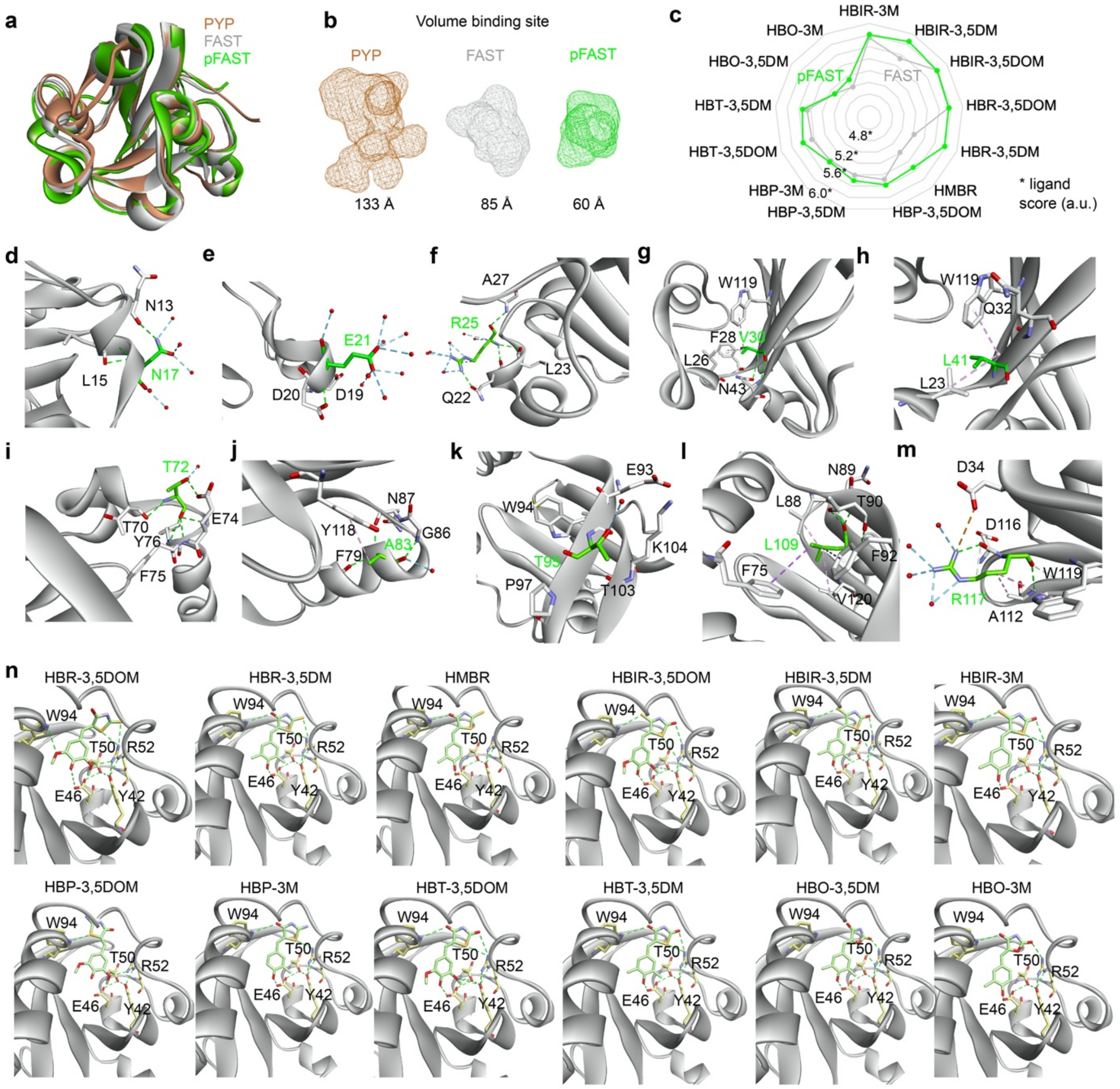


**Fig. S6. Structural homology models. (a)** Structural models of FAST and pFAST generated from the tridimensional crystal structure of the *Halorhodospira halophila* Photoactive Yellow Protein PYP (PDB: 6P4I). **(b)** Volumes of the ligand binding sites of PYP, FAST and pFAST. **(c)** In silico binding affinities (ligscore 2) of FAST and pFAST for the HBIR, HBR, HBR, HBT and HBO series. **(d-m)** Mapping of the residues involved in polar (green dashed lines) and apolar interactions (purple dashed lines) with the residues found mutated in pFAST. **(n)** Molecular docking of pFAST-chromophore assemblies.

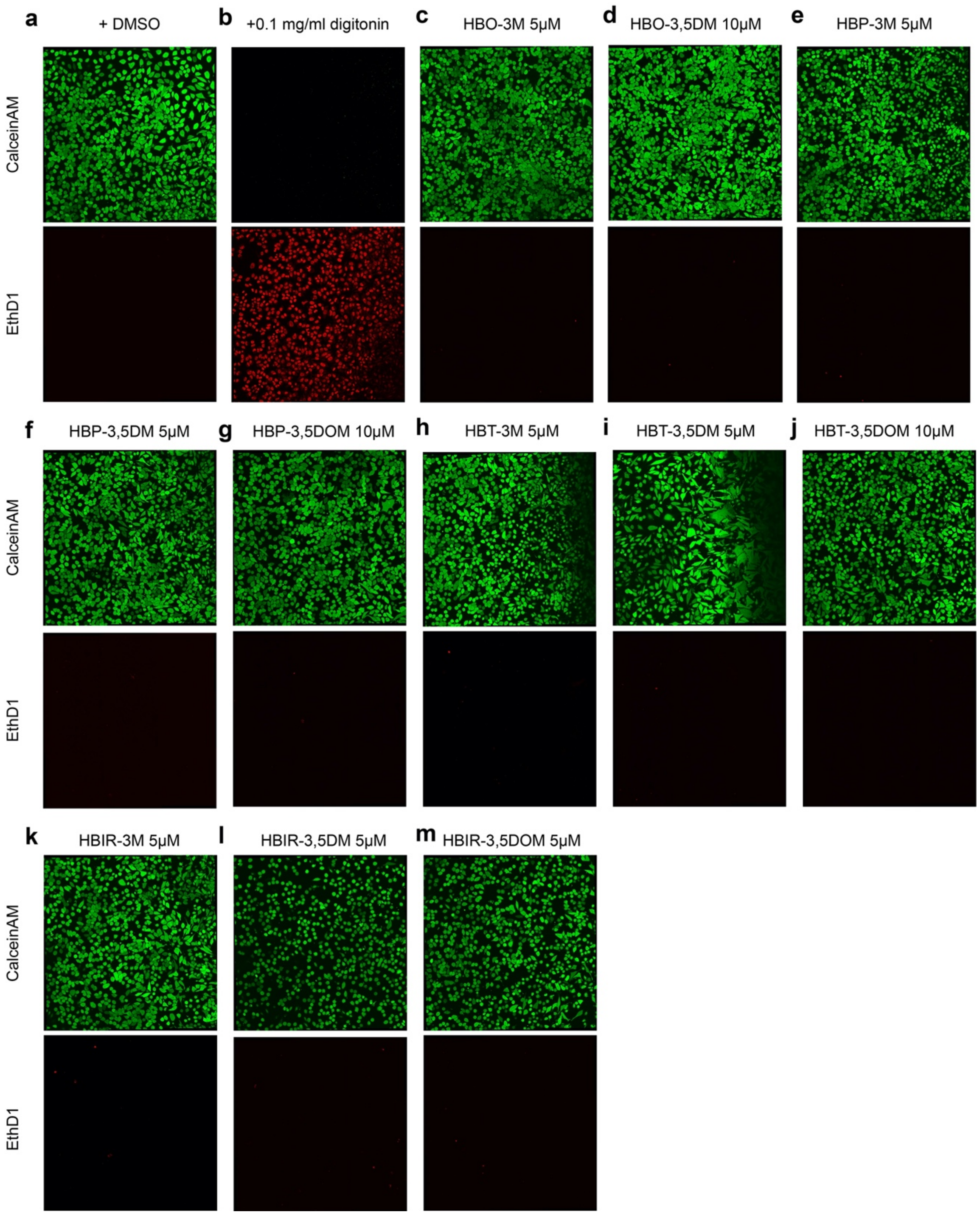


**Fig. S7. Two-color fluorescence viability assay.** HeLa cells were incubated for 24 h with solutions of **(a-b)** 0.1 % of DMSO, **(c)** HBO-3M at 5 μM, **(d)** HBO-3,5DM at 10 μM, **(e)** HBP-3M at 5 μM, **(f)** HBP-3,5DM at 5 μM, **(g)** HBP-3,5DOM at 10 μM, **(h)** HBT-3M at 5 μM, **(i)** HBT-3,5DM at 5 μM, **(j)** HBT-3,5DOM at 5 μM , **(k)** HBIR-3M at 5 μM, **(l)** HBIR-3,5DM at 5 μM and **(m)** HBIR-3,5DOM at 5 μM. Control experiments of HeLa cells non-incubated with dye (**a**, live cells) or incubated for 30 min with 0.1 mg/ml digitonin (**b**, dead cells) are shown. Cell viability was tested with calceinAM and EthD1 probes (LIVE/DEAD® viability/cytotoxicity assay kit). CalceinAM is a cell-permeant profluorophore cleaved by intracellular esterases releasing a uniform green fluorescence polyanionic calcein in live cells (green channel). EthD1 (Ethidium homodimer-1) is a non-permeant nucleic acid fluorescent stain that enters only cells with damaged membranes and undergoes a fluorescence enhancement upon binding to nucleic acids, thereby producing a bright red fluorescence in dead cells (red channel). Cell fluorescence was evaluated by confocal microscopy. Identical imaging settings were used for the different experiments. The experiment shows that none of the chromophores are toxic for HeLa cells at the concentrations used for imaging.

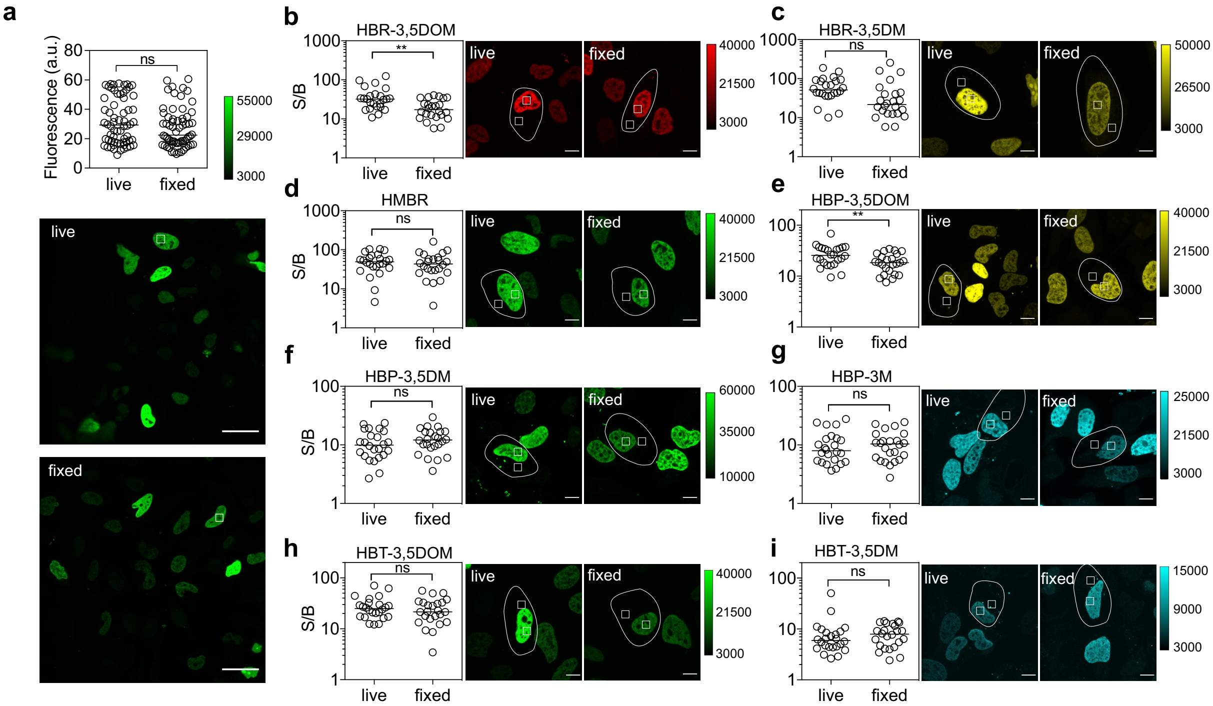


**Fig. S8. Imaging of H2B-pFAST in live and fixed HeLa cells after addition of various fluorogenic chromophores. (a)** Nuclear fluorescence in live and fixed cells incubated with 5 µM of HMBR and imaged with identical microscope settings (n = 64 cells, 2 experiments). Median values are reported and a Wilcoxon test was conducted to compare live and fixed cell populations (ns = not significative). Scale bars, 40 µm. **(b-i)** Comparison of signal (nucleus) to background (cytosol) (S/B) ratio between live and fixed cells incubated with **(b)** HBR-3,5DOM at 5 µM, **(c)** HBR-3,5DM at 5 µM, **(d)** HMBR at 5 µM, **(e)** HBP-3,5DOM at 10 µM, **(f)** HBP-3,5DM at 5 µM, **(g)** HBP-3M at 5 µM, **(h)** HBT-3,5DOM at 5 µM and **(i)** HBT-3,5DM at 5 µM. For each experiment, live and fixed cells were imaged with identical microscope settings (n = 24 cells, 2 experiments). Median of S/B ratios are reported and a Wilcoxon test was conducted to compare live and fixed cell populations (ns = not significative, * $p < 0.05$ and ** $p < 0.01$). (see **Table S13** for imaging settings). Scale bars, 10 µm.

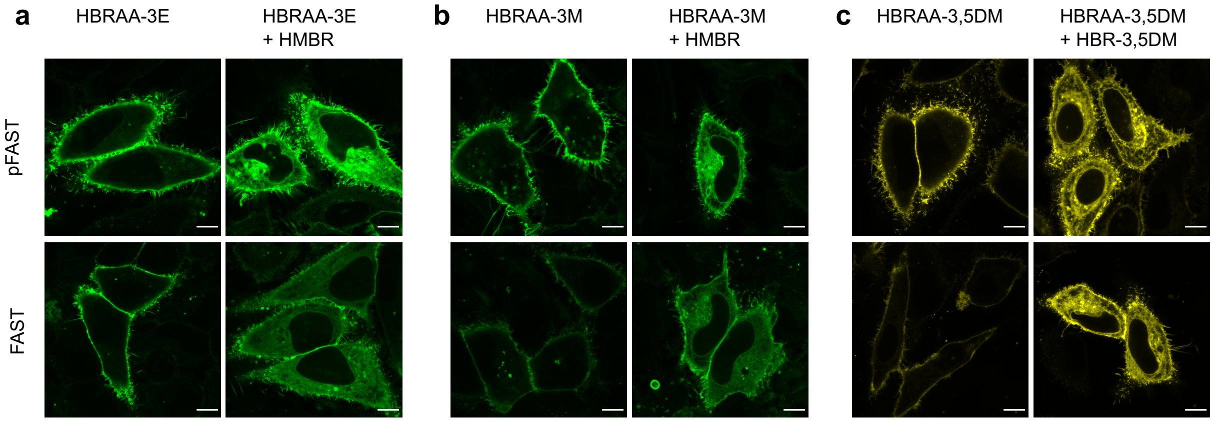


**Fig. S9. Selective imaging of cell-surface proteins.** Confocal micrographs of HeLa cells expressing a secreted transmembrane domain fused to pFAST versus FAST labeled with impermeant fluorogens **(a)** HBRAA-3E at 10 µM, **(b)** HBRAA-3M at 10 µM and **(c)** HBRAA-3,5DM at 10 µM. Subsequent addition of 5 µM of membrane-permeant HMBR **(a,b)** or HBR-3,5DM **(c)** revealed the total pool of proteins expressed at the surface and within the secretory pathway. For each experiment, FAST and pFAST were imaged under the same imaging settings. The detection settings were adjusted to take into account the difference of brightness of impermeant and membrane-permeant fluorogens for each experiment. (see **Table S13** for imaging settings). Scale bars, 10 µm.

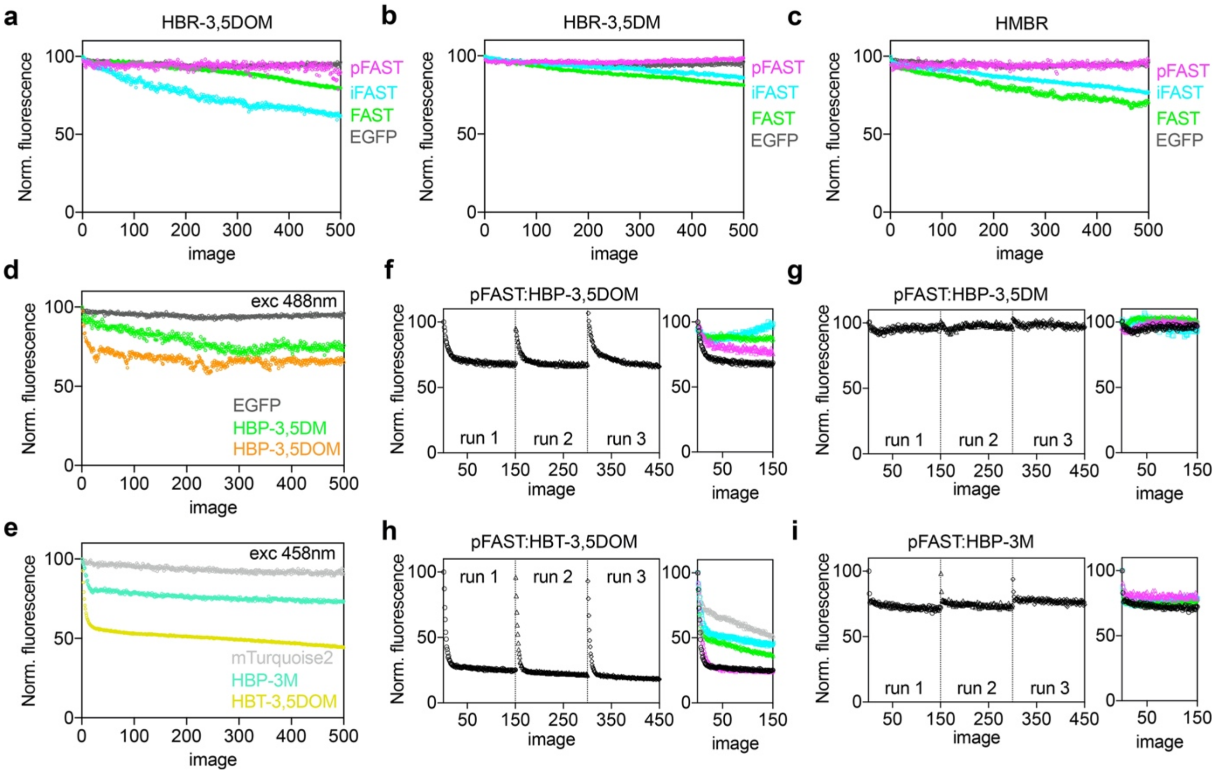


**Fig. S10**. **In-cell photostability of pFAST. (a-c)** Photostability of pFAST compared to FAST and iFAST in presence of 10 μM HBR-3,5DOM (a), 5 μM HBR-3,5DM (b) and 5 μM HMBR (c). Photostability of EGFP is also given for comparison. Cells were illuminated with a 488 nm laser excitation (with a power of 4.4 $kW/cm^2$ at the specimen plane) and 500 images were acquired every 2 s, n = 3 cells per curve. **(d)** Photostability of pFAST in presence of 10 μM HBP-3,5DOM and 5 μM HBP-3,5DM under 488 nm laser excitation (with a power of 4.4 $kW/cm^2$ at the specimen plane). Photostability of EGFP is also given for comparison. 500 images were acquired every 2 s, n = 3 cells per curve. **(e)** Photostability of pFAST in presence of 5 μM HBT-3,5DOM and 5 μM HBP-3M under continuous 458 nm laser excitation (with a power of 3.5 $kW/cm^2$ at the specimen plane). Photostability of mTurquoise2 is also given for comparison. 500 images were acquired every 2 s, n = 3 cells per curve. **(f-g)** Photostability of pFAST in presence of 10 μM HBP-3,5DOM (f) and 5 μM HBP-3,5DM (g) under 488 nm laser excitation with a power of 4.7 $kW/cm^2$ (black and green curves) and 2.4 $kW/cm^2$ (pink and cyan curves) at the specimen plan. 150 images were acquired every 2s (black and pink curves) and 5s (green and cyan curves) followed by 60s in the dark before acquisition was restarted. **(h-i)** Photostability of pFAST in presence of 5 μM HBT-3,5DOM (h) and 5 μM HBP-3M (i) under 458 nm laser excitation with a power of 3.7 $kW/cm^2$ (black and green curves) and 2.4 $kW/cm^2$ (pink, cyan and gray curves) at the specimen plan. 150 images were acquired every 2 s (black and pink curves), every 5 s (green and cyan curves) and every 10 s (gray curve only for HBT-3,5DOM) followed by 60 s in the dark before acquisition was restarted. **(a-i)** The proteins were expressed in the cytoplasm in HeLa cells and images were acquired using a scanning confocal microscope with a pixel dwell time of 2.55 μs (see **Table S13** for imaging settings).

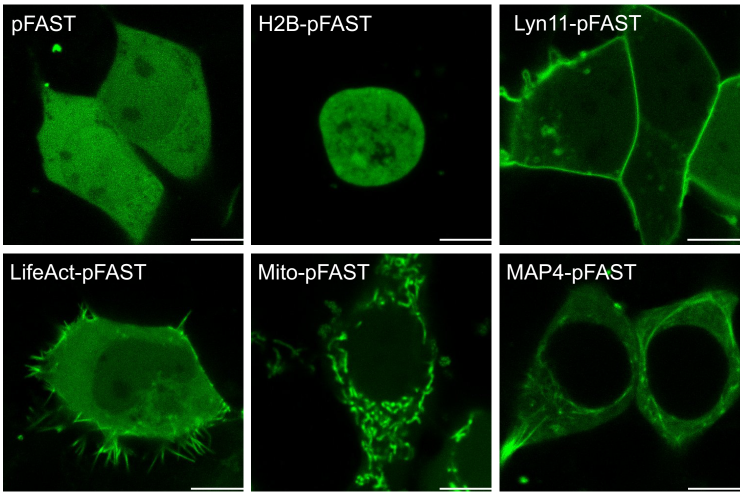


**Fig. S11.** Selective imaging of pFAST in live mammalian cells. Confocal micrographs of live HEK293T cells expressing pFAST fused to: histone H2B, lyn11 (inner membrane-targeting motif), LifeAct (actin binding peptide domain), mito (mitochondrial targeting motif) and to microtubule-associated protein (MAP) 4 and labeled with 5 μM HBP-3,5DM (see **Table S13** for imaging settings). Scale bars, 10 μm.

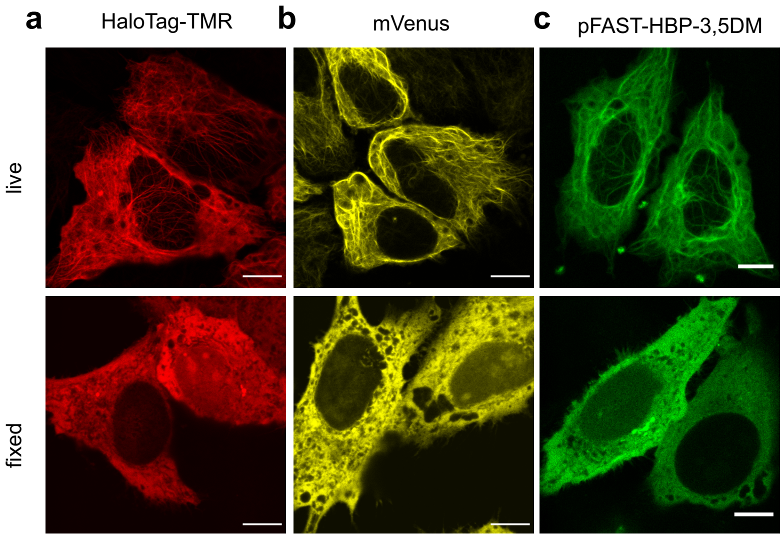


**Fig. S12**. Confocal micrographs of live and fixed HeLa cells expressing MAP4 fused to **(a)** HaloTag labeled with 2.5 μM of HaloTag® TMR Ligand, **(b)** mVenus and **(c)** pFAST labeled with HBP-3,5DM at 5 μM (see **Table S13** for imaging settings). Scale bars, 10 μm.

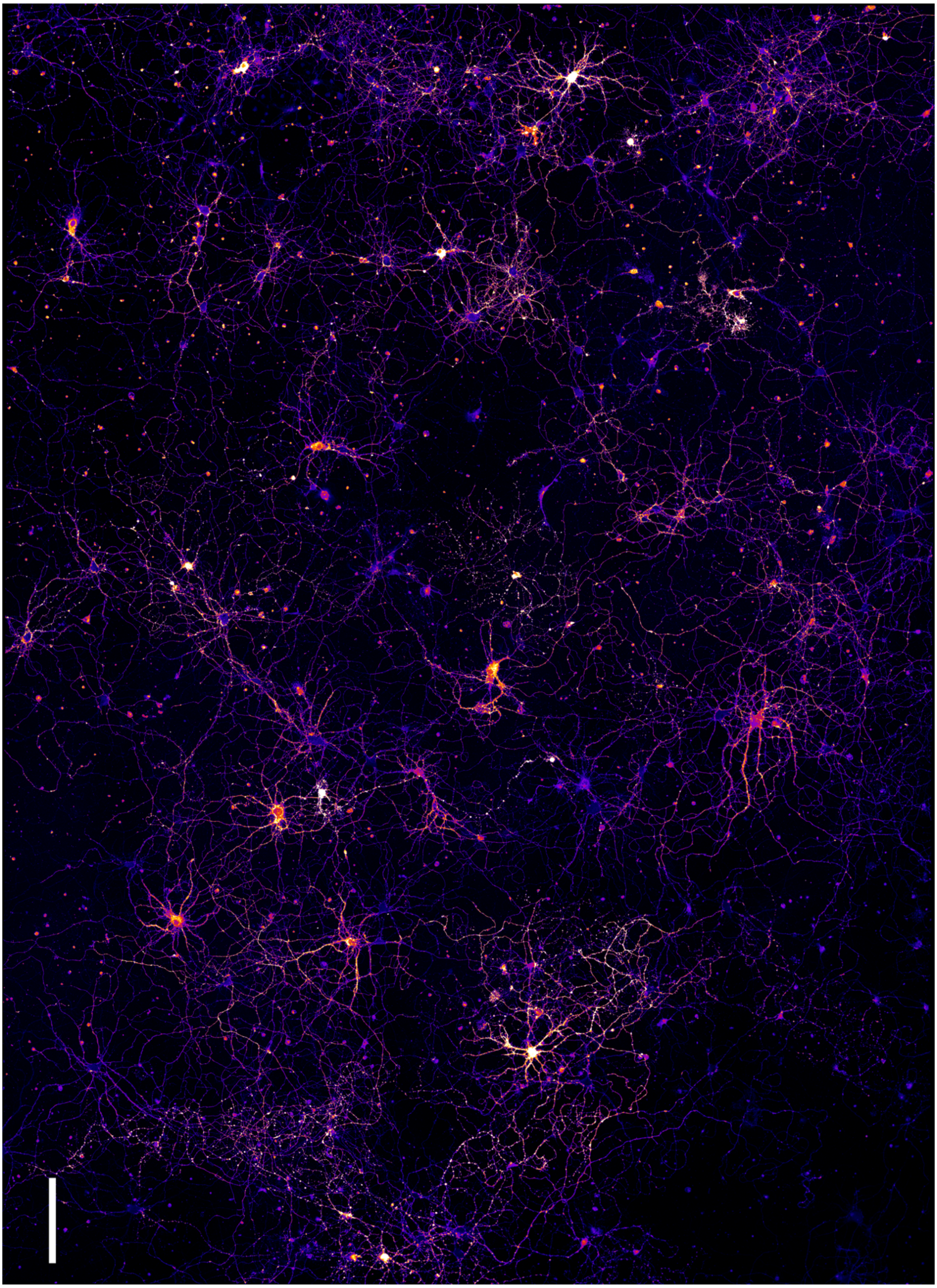

**Fig. S13. Selective imaging of hippocampal neuronal network.** Tiles of confocal micrographs of dissociated hippocampal neurons transfected with a plasmid encoding pFAST fused to lyn11 (inner membrane targeting motif), and labeled with 10 μM HBR-3,5DOM (see **Table S13** for imaging settings). The huge field of view enables to illustrate the high transfection rate and viability of transfected fragile cells like neurons. Artificial look up table 'fire' color is used to illustrate intensity dynamics. Scale bar, 200 μm.

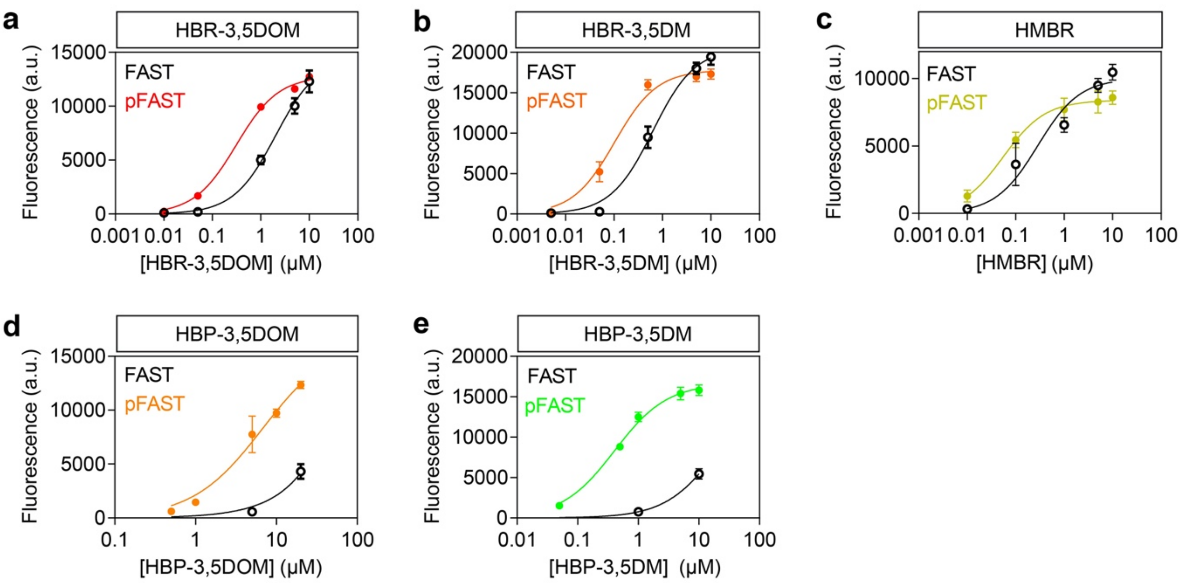


**Fig. S14**. **Labeling efficiency in live cells.** Fluorescence of HEK 293T cells expressing cytoplasmic pFAST (colored lines) versus FAST (black lines) in the presence of increased concentrations of **(a)** HBR-3,5DOM, **(b)** HBR-3,5DM, **(c)** HMBR, **(d)** HBP-3,5DOM and **(e)** HBP-3,5DM (n = 3 replicates). Fluorescence was measured by flow cytometry.

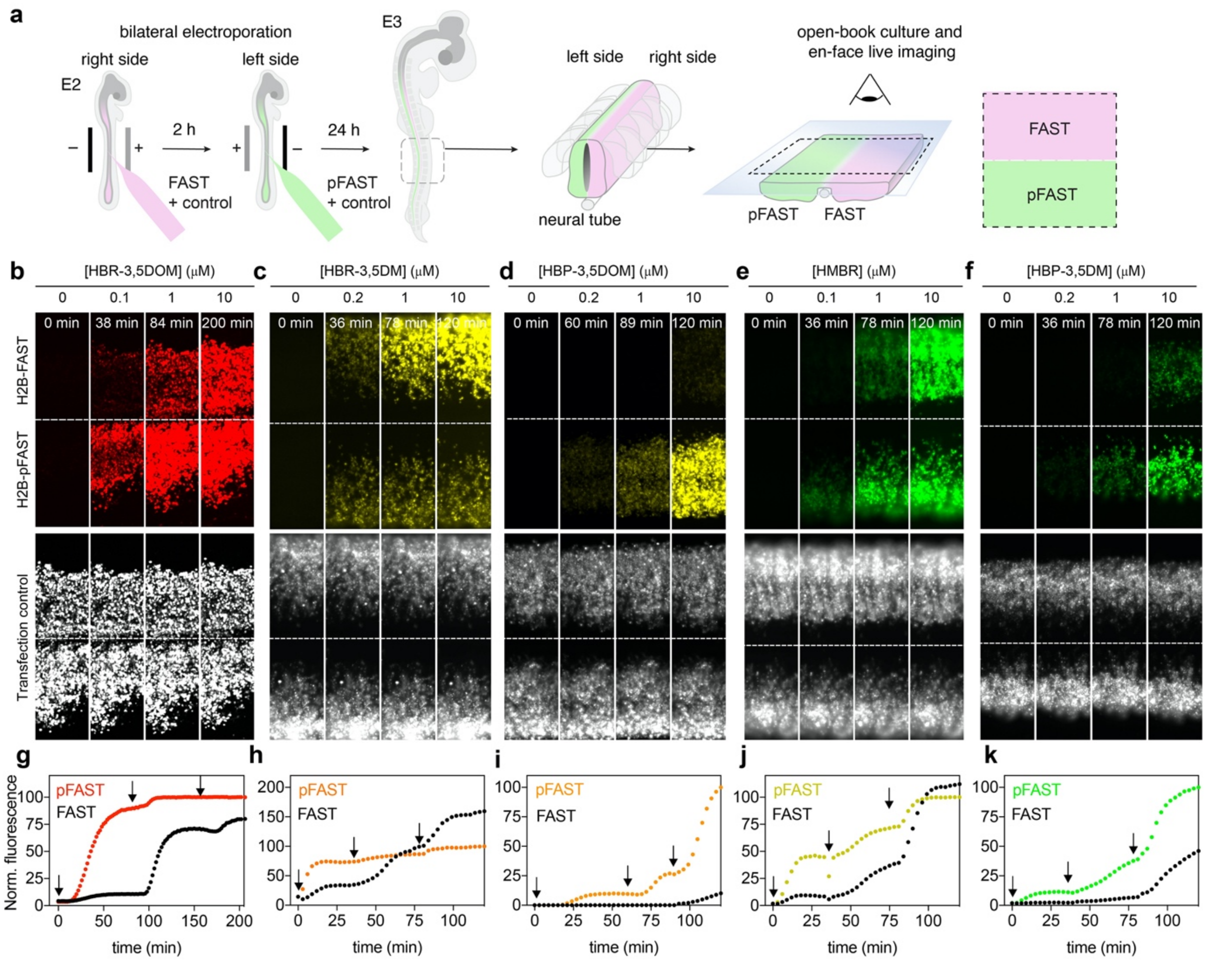


**Fig. S15**. **Labeling efficiency in chicken embryos. (a-f)** Plasmids encoding H2B-pFAST and H2B-FAST were electroporated in each side of the neural tube in ovo at embryonic day 2 (E2, HH stage 13-14). (b) EGFP or (c-f) mCherry reporters were co-injected with each construct as a transfection efficiency control. 24 h later, embryos with homogeneous bilateral reporter expression in the neural tube were dissected and imaged. Time-lapse imaging upon sequential addition of fluorogenic chromophore were acquired using (b) spinning-disk confocal or (c-f) widefield fluorescence microscopy. Conditions: **(b)** 0.1, 1 and 10 μM HBR-3,5DOM (see also **Movie S1)** ; **(c)** 0.2, 1 and 10 μM HBR-3,5DM ; **(d)** 0.2, 1 and 10 μM of HBP-3,5DOM ; **(e)** 0.1, 1 and 10 μM of HMBR and **(f)** 0.2, 1 and 10 μM of HBP-3,5DM (see **Table S13** for imaging settings). **(g-k)** Fluorescence intensities of pFAST and FAST were analyzed over time and normalized by the maximal fluorescence value of pFAST for each fluorogenic chromophore. The sequential addition of the fluorogenic chromophore at the different concentrations are indicated by arrows.

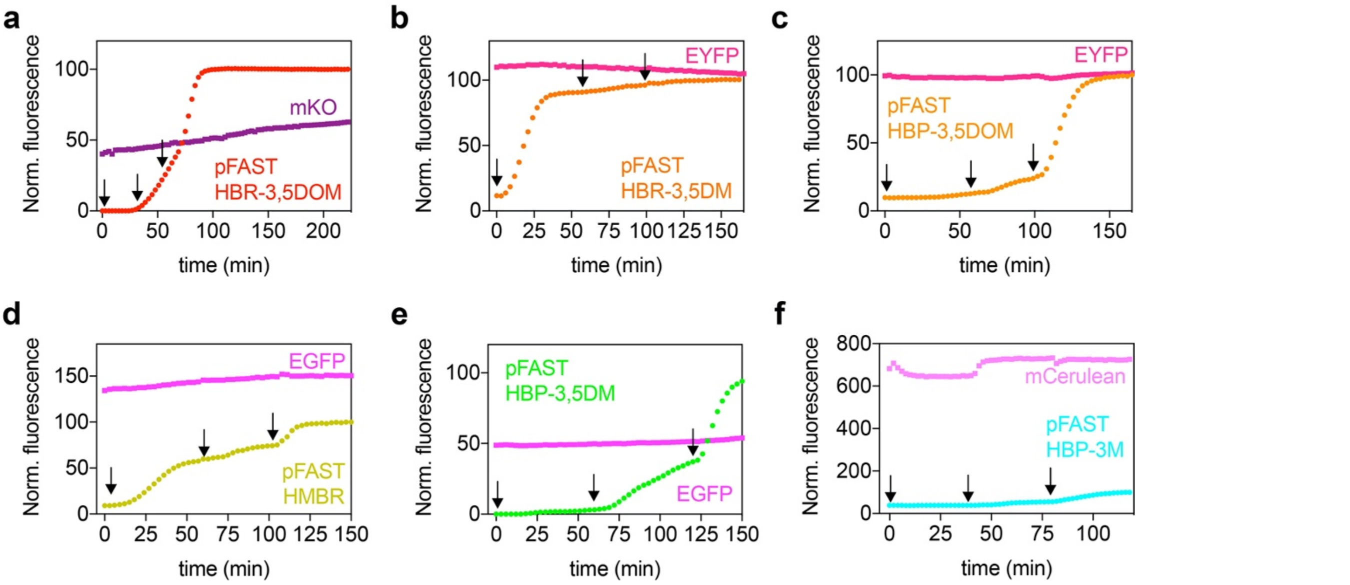


**Fig. S16. Comparison of pFAST with various fluorescent proteins in chicken embryos**. Embryos expressing pFAST and the fluorescent proteins (a) mKO, (b,c) EYFP, (d,e) EGFP and (f) mCerulean in each side of the neural tube were dissected and labeled by sequential additions of various concentrations of fluorogenic chromophores (arrows indicate additions). **(a)** 0.1, 1, 10 μM of HBR-3,5DOM; **(b)** 0.2, 1, 10 μM of HBR-3,5DM; **(c)** 0.2, 1, 10 μM of HBP-3,5DOM; **(d)** 0.2, 1, 10 μM of HMBR; **(e)** 0.2, 1, 10 μM of HBP-3,5DM and **(f)** 0.2, 1, 10 μM of HBP-3M. Time-lapse (a) spinning-disk confocal or (b-f) widefield fluorescence imaging allowed to monitor the temporal evolution of the fluorescence intensity (**Fig. 5** in main text shows the initial and final images). Fluorescence intensities were normalized by the maximal fluorescence value of pFAST for each experiment.

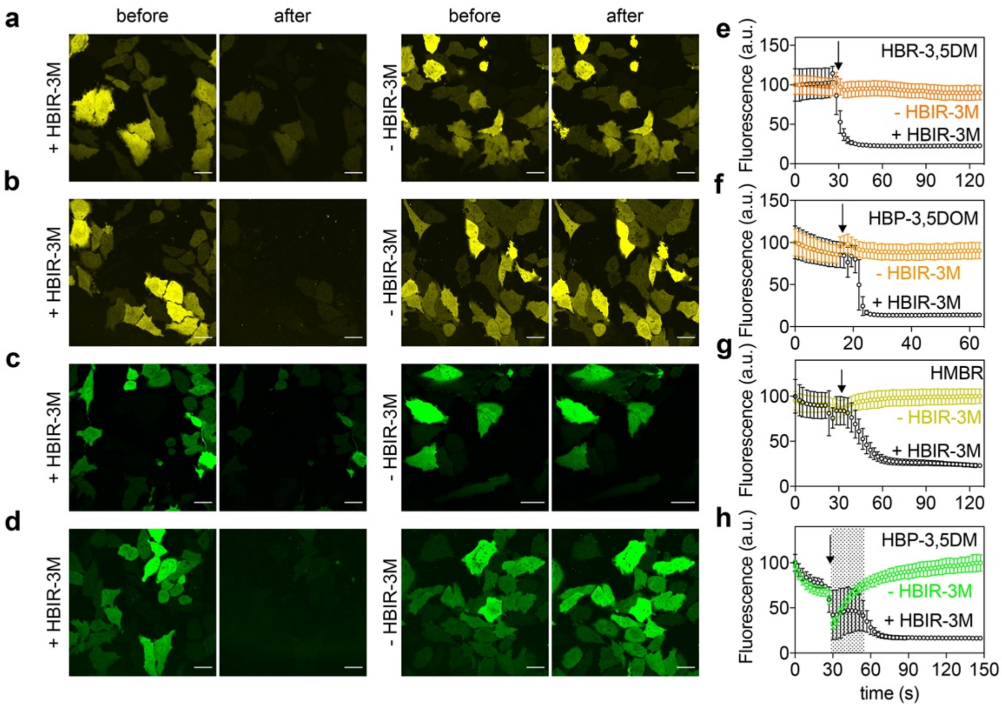


**Fig. S17. Reversible labeling of pFAST in live cells by chromophore replacement. (a-d)** Confocal micrographs of HeLa cells expressing cytoplasmic pFAST - initially labeled with (a) 1 μM HBR-3,5DM, (b) 10 μM HBP-3,5DOM, (c) 1 μM HMBR and (d) 5 μM HBP-3,5DM - acquired before and after addition of 10 μM HBIR-3M dark-competitor (the concentration of fluorogenic chromophore was kept constant during the overall experiment) (see **Table S13** for imaging settings). Scale bars, 10 μm. **(e-h)** Temporal evolution of fluorescence intensities upon addition of HBIR-3M (or mock solutions) as indicated by arrows (n = 3 cells) **(**see also **Movie S5).** Note that the artefactual jump of fluorescence on (h) (grey area) was due to a change of focus upon addition of the HBIR-3M solution or the mock solution.

**a**

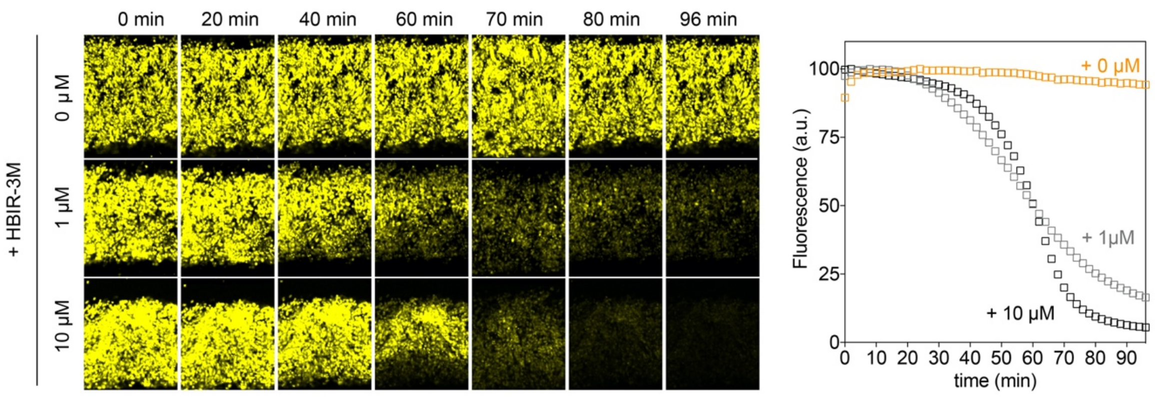


**b**

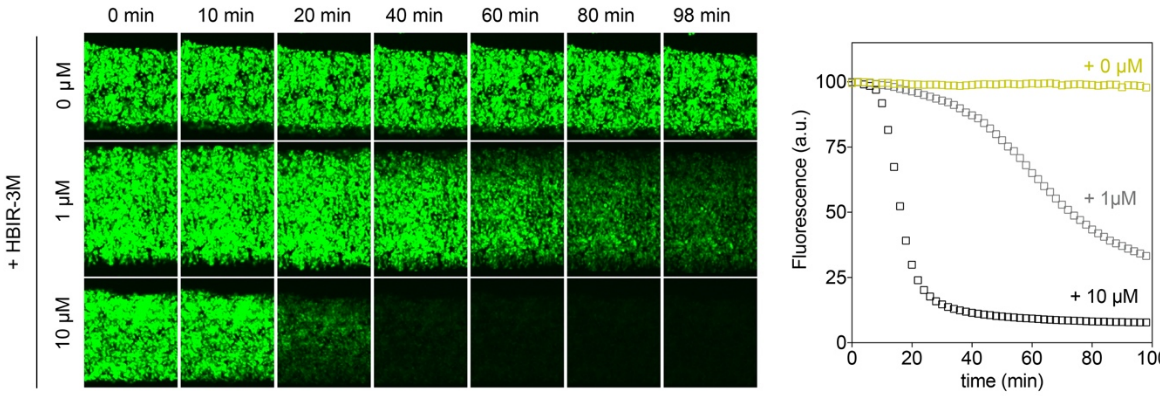


**Fig. S18. Reversible labeling of pFAST in chicken embryos by chromophore replacement.** Embryos expressing H2B-pFAST in the neural tube were dissected and labeled with **(a)** 5 μM of HBP-3,5DOM and with **(b)** 1 μM of HMBR for 40 minutes (see also **Movie S6**). The labeling solution was then removed, the samples were washed once with PBS and fresh medium supplemented with 0 μM, 1 μM and 10 μM of the dark competitor HBIR-3M was added. Time-lapse spinning-disk confocal imaging allowed to monitor the temporal evolution of the fluorescence intensity after addition of the dark competitor (see **Table S13** for imaging settings). The graphs show the temporal evolution of the fluorescence intensities.

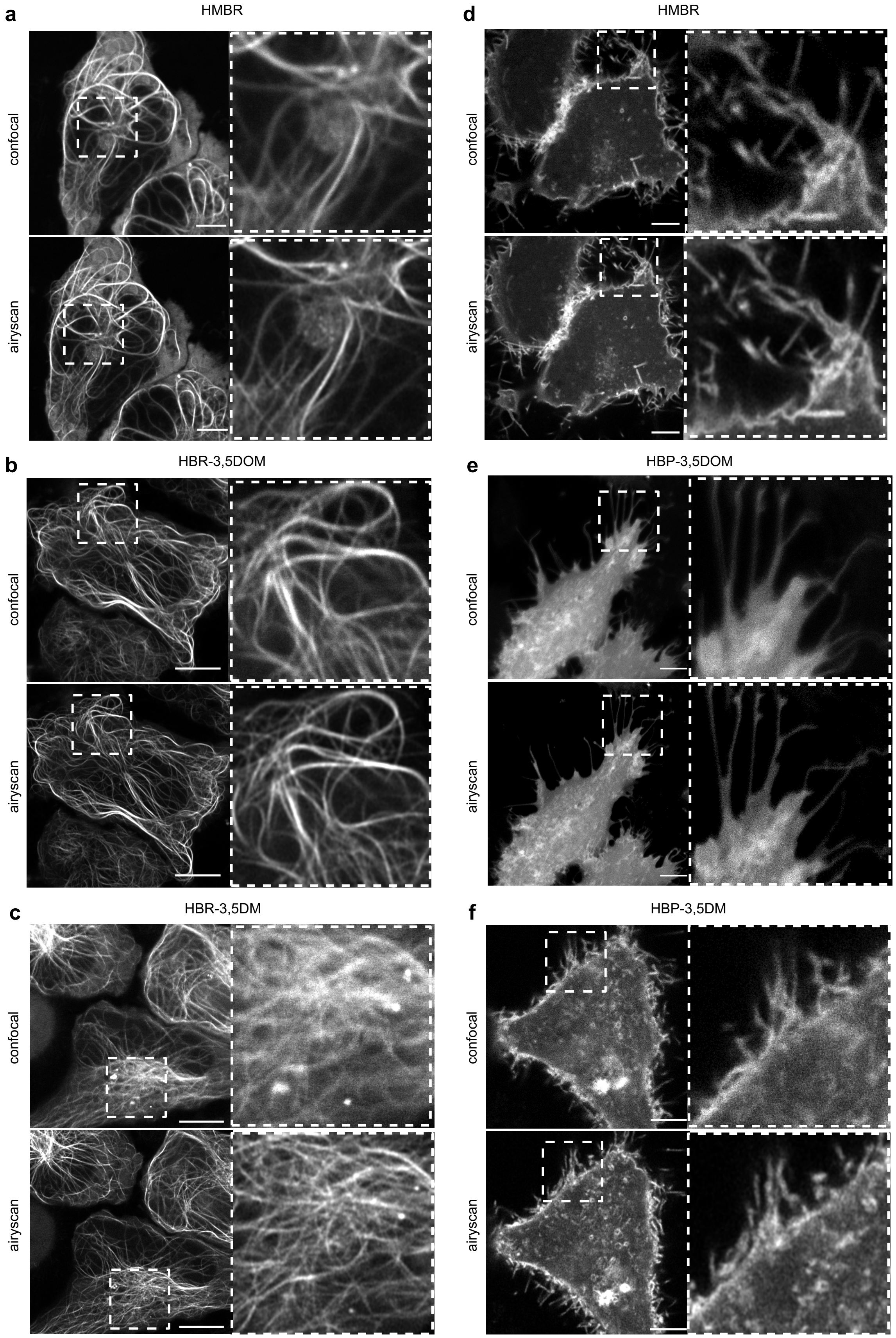


**Fig. S19. Airyscan confocal imaging of live mammalian cells expressing (a-c)** MAP4-pFAST (microtubule associated protein) fusion or **(d-f)** Lyn11-pFAST (membrane targeting motif) fusion and labeled with (a,d) 5 μM HMBR, (b) 5 μM HBR-3,5DOM, (c) 5 μM HBR-3,5DM, (e) 10 μM HBP-3,5DOM and (f) 5 μM HBP-3,5DM (see **Table S13** for imaging settings). Scale bars, 10 μm.

**Table S1.** Properties of FAST with various chromophores in PBS pH 7.4.

| Chromophore | $\lambda_{abs}$ (nm) | $\Delta\lambda_{abs}$ (nm) | $\lambda_{em}$ (nm) | $\varepsilon$ ($mM^{-1}cm^{-1}$) | $\phi$ | Molecular brightness | $K_D$ (µM) |
|---|---|---|---|---|---|---|---|
| HBR-3,5DOM (ref 2) | 520 | 114 | 600 | 39 | 0.31 | 12,000 | 0.97 |
| HBR-3,5DM (ref 2) | 499 | 97 | 562 | 48 | 0.49 | 24,000 | 0.08 |
| HMBR (ref 2) | 481 | 79 | 540 | 44 | 0.23 | 10,000 | 0.13 |
| HBP-3,5DOM | 488 | 121 | 556 | 31 | 0.29 | 9,000 | ~100 |
| HBP-3,5DM | 464 | 105 | 522 | 30 | 0.16 | 4,000 | 3.7 |
| HBP-3M | 447 | 87 | 506 | 21 | 0.044 | 800 | 6.9 |
| HBT-3,5DOM | 449 | 92 | 532 | 54 | 0.22 | 12,000 | 3.3 |
| HBT-3,5DM | 434 | 82 | 500 | 25 | 0.072 | 2,000 | 1.7 |
| HBT-3M | 415 | - | 480 | 32 | 0.006 | 200 | 0.24 |
| HBO-3,5DM | 409 | 80 | 481 | 4 | 0.20 | 700 | 32 |
| HBO-3M | 394 | 68 | 471 | 11 | 0.056 | 600 | 6.4 |

Abbreviations are as follows: $\lambda_{abs}$ wavelength of maximal absorption; $\Delta\lambda_{abs} = \lambda_{abs,bound} - \lambda_{abs,unbound}$ absorption red-shift upon chromophore binding; $\lambda_{em}$ wavelength of maximal emission; $\varepsilon$, molar absorptivity at $\lambda_{abs}$ (standard error is typically 10%); $\phi$, fluorescence quantum yield; molecular brightness = $\phi \times \varepsilon$; $K_D$ thermodynamic dissociation constant.

**Table S2.** Properties of oFAST with various chromophores in PBS pH 7.4

| Chromophore | $\lambda_{abs}$ (nm) | $\Delta\lambda_{abs}$ (nm) | $\lambda_{em}$ (nm) | $\varepsilon$ ($mM^{-1}cm^{-1}$) | $\phi$ | Molecular brightness | $K_D$ (µM) |
|---|---|---|---|---|---|---|---|
| HBR-3,5DOM | 519 | 115 | 600 | 44 | 0.33 | 14,000 | 0.1 |
| HBR-3,5DM | 500 | 98 | 560 | 53 | 0.38 | 20,000 | 0.01 |
| HMBR | 481 | 79 | 540 | 47 | 0.24 | 11,000 | 0.19 |
| HBP-3,5DOM | 489 | 122 | 556 | 28 | 0.30 | 8,000 | 3.0 |
| HBP-3,5DM | 465 | 104 | 519 | 33 | 0.21 | 7,000 | 0.20 |
| HBP-3M | 445 | 85 | 505 | 40 | 0.054 | 2,000 | 0.37 |
| HBT-3,5DOM | 449 | 92 | 533 | 68 | 0.21 | 15,000 | 0.30 |
| HBT-3,5DM | 435 | 83 | 501 | 31 | 0.090 | 3,000 | 0.16 |
| HBO-3,5DM | 411 | 82 | 482 | 6 | 0.30 | 2,000 | 3.0 |
| HBO-3M | 394 | 68 | 470 | 15 | 0.11 | 1,600 | 0.73 |

Abbreviations are as follows: $\lambda_{abs}$ wavelength of maximal absorption; $\Delta\lambda_{abs} = \lambda_{abs,bound} - \lambda_{abs,unbound}$ absorption red-shift upon chromophore binding; $\lambda_{em}$ wavelength of maximal emission; $\varepsilon$ molar absorptivity at $\lambda_{abs}$ (standard error is typically 10%); $\phi$ fluorescence quantum yield; molecular brightness = $\phi \times \varepsilon$; $K_D$ thermodynamic dissociation constant.

**Table S3.** Properties of tFAST with various chromophores in PBS pH 7.4.

| Chromophore | $\lambda_{abs}$ (nm) | $\Delta\lambda_{abs}$ (nm) | $\lambda_{em}$ (nm) | $\varepsilon$ ($mM^{-1}cm^{-1}$) | $\phi$ | Molecular brightness | $K_D$ (µM) |
|---|---|---|---|---|---|---|---|
| HBR-3,5DOM | 520 | 115 | 600 | 46 | 0.33 | 15,000 | 0.07 |
| HBR-3,5DM | 501 | 99 | 561 | 56 | 0.44 | 25,000 | 0.01 |
| HMBR | 481 | 79 | 541 | 52 | 0.24 | 12,000 | 0.19 |
| HBP-3,5DOM | 488 | 121 | 555 | 28 | 0.30 | 8,000 | 4.0 |
| HBP-3,5DM | 464 | 104 | 522 | 33 | 0.25 | 8,000 | 0.36 |
| HBP-3M | 447 | 87 | 505 | 39 | 0.075 | 3,000 | 0.56 |
| HBT-3,5DOM | 449 | 92 | 532 | 60 | 0.24 | 15,000 | 0.44 |
| HBT-3,5DM | 435 | 83 | 497 | 30 | 0.11 | 3,000 | 0.33 |
| HBO-3,5DM | 410 | 81 | 485 | 6 | 0.22 | 1,000 | 5.0 |
| HBO-3M | 393 | 67 | 471 | 13 | 0.069 | 1,000 | 0.8 |

Abbreviations are as follows: $\lambda_{abs}$ wavelength of maximal absorption; $\Delta\lambda_{abs} = \lambda_{abs,bound} - \lambda_{abs,unbound}$ absorption red-shift upon chromophore binding; $\lambda_{em}$ wavelength of maximal emission; $\varepsilon$ molar absorptivity at $\lambda_{abs}$ (standard error is typically 10%); $\phi$ fluorescence quantum yield; molecular brightness = $\phi \times \varepsilon$; $K_D$ thermodynamic dissociation constant.

**Table S4.** Properties of pFAST with various chromophores in PBS pH 7.4.

| Chromophore | $\lambda_{abs}$ (nm) | $\Delta\lambda_{abs}$ (nm) | $\lambda_{em}$ (nm) | $\varepsilon$ ($mM^{-1}cm^{-1}$) | $\phi$ | Molecular brightness | $K_D$ ( $\mu$M) |
|---|---|---|---|---|---|---|---|
| HBIR-3,5DOM | 562 | 128 | 616 | 40 | 0.10 | 3,000 | 0.04 |
| HBIR-3,5DM | 536 | 106 | 578 | 25 | 0.015 | 400 | 0.07 |
| HBIR-3M | 514 | 84 | 567 | 12 | 0.003 | 50 | 0.005 |
| HBR-3,5DOM | 520 | 115 | 600 | 44 | 0.35 | 15,000 | 0.06 |
| HBR-3,5DM | 501 | 99 | 561 | 49 | 0.44 | 22,000 | 0.01 |
| HMBR | 481 | 79 | 542 | 54 | 0.23 | 13,000 | 0.01 |
| HBRAA-3,5DM | 524 | 118 | 578 | 58 | 0.22 | 12,000 | 1.8 |
| HBRAA-3E | 506 | 96 | 558 | 53 | 0.05 | 3,000 | 0.05 |
| HBRAA-3M | 502 | 96 | 554 | 64 | 0.08 | 5,000 | 0.23 |
| HBP-3,5DOM | 487 | 120 | 554 | 37 | 0.33 | 12,000 | 1.9 |
| HBP-3,5DM | 465 | 105 | 520 | 37 | 0.27 | 10,000 | 0.15 |
| HBP-3M | 447 | 87 | 503 | 35 | 0.10 | 4,000 | 0.22 |
| HBT-3,5DOM | 449 | 92 | 532 | 60 | 0.27 | 16,000 | 0.20 |
| HBT-3,5DM | 433 | 81 | 499 | 33 | 0.10 | 3,000 | 0.17 |
| HBO-3,5DM | 411 | 82 | 483 | 6 | 0.23 | 1,000 | 2.4 |
| HBO-3M | 392 | 66 | 473 | 14 | 0.076 | 1,000 | 0.43 |

Abbreviations are as follows: $\lambda_{abs}$ wavelength of maximal absorption; $\Delta\lambda_{abs} = \lambda_{abs,bound} - \lambda_{abs,unbound}$ absorption red-shift upon chromophore binding; $\lambda_{em}$ wavelength of maximal emission; $\varepsilon$, molar absorptivity at $\lambda_{abs}$ (standard error is typically 10%); $\phi$ fluorescence quantum yield; molecular brightness = $\phi \times \varepsilon$; $K_D$ thermodynamic dissociation constant.

**Table S5.** Properties of the mutants isolated from the selection with HBO-3M compared to FAST in PBS pH 7.4.

| Clone | Mutations (relative to FAST) | $\lambda_{abs}$ (nm) | $\lambda_{em}$ (nm) | $\varepsilon$ ($mM^{-1}cm^{-1}$) | $\phi$ | Molecular brightness | $K_D$ ($\mu$M) |
|---|---|---|---|---|---|---|---|
| FAST | | 394 | 471 | 11 | 0.056 | 600 | 6.4 |
| A-R6.8 | D65V / E93D / M109L / S117R | 395 | 468 | 15 | 0.090 | 1,300 | 2.6 |
| A-R6.11 | V83I / M109L | 394 | 468 | 15 | 0.074 | 1,100 | 1.4 |
| A-R7.1 | M109L | 394 | 470 | 13 | 0.081 | 1,000 | 4.0 |
| A-R7.2 | Q41L / M109L | 394 | 468 | 12 | 0.092 | 1,000 | 1.1 |
| A-R7.10 | V83I / T103I | 394 | 468 | 12 | 0.088 | 1,000 | 4.4 |
| A-R7.12 | K17R / A30V / E74K / E93V | 386 | 466 | 13 | 0.097 | 1,200 | 2.0 |
| A-R7.14 | T50S / E93Q / M95I | 382 | 468 | 13 | 0.077 | 1,000 | 1.4 |

Abbreviations are as follows: $\lambda_{abs}$ wavelength of maximal absorption; $\lambda_{em}$ wavelength of maximal emission; $\varepsilon$ molar absorptivity at $\lambda_{abs}$ (standard error is typically 10%); $\phi$ fluorescence quantum yield; molecular brightness = $\phi \times \varepsilon$; $K_D$ thermodynamic dissociation constant.

**Table S6.** Properties of the mutants isolated from the selection with HBO-3,5DM compared to FAST in PBS pH 7.4.

| Clone | Mutations (relative to FAST) | $\lambda_{abs}$ (nm) | $\lambda_{em}$ (nm) | $\varepsilon$ ($mM^{-1}cm^{-1}$) | $\phi$ | Molecular brightness | $K_D$ ($\mu$M) |
|---|---|---|---|---|---|---|---|
| FAST | | 409 | 481 | 4 | 0.20 | 800 | 32 |
| A-R6.6 | V83I / M109L | 402 | 481 | 5 | 0.22 | 1,200 | 11 |
| A-R6.18 | H3Q / D48A / T103I / M109L | 408 | 479 | 5 | 0.22 | 1,000 | 6.2 |
| A-R7.1 | N13I / D48E / D71N / V83G / M95V | 408 | 483 | 5 | 0.24 | 1,100 | 5.0 |
| A-R7.2 | Q41L / M95I / M109L | 406 | 481 | 6 | 0.25 | 1,500 | 6.1 |
| A-R7.5 | K60R / V83I / K104R / S117R | 406 | 481 | 5 | 0.22 | 1,000 | 7.1 |
| A-R7.7 | D71N / M109L / S117R | 407 | 482 | 5 | 0.25 | 1,200 | 4.4 |
| A-R7.7-1 | D71N / M109L / S117R + Q41L | 413 | 480 | 5 | 0.26 | 1,300 | 2.5 |
| A-R7.7-2 | D71N / M109L / S117R + Q41L + D48E | 411 | 480 | 6 | 0.27 | 1,500 | 2.1 |
| A-R7.7-3 | D71N / M109L / S117R + Q41L + D65V | 411 | 483 | 7 | 0.27 | 1,800 | 2.4 |
| A-R7.7-4 = oFAST | D71N / M109L / S117R + Q41L + V83I | 411 | 482 | 6 | 0.30 | 2,000 | 3.0 |
| A-R7.7-5 | D71N / M109L / S117R + Q41L + M95I | 411 | 482 | 5 | 0.26 | 1,400 | 3.0 |

Abbreviations are as follows: $\lambda_{abs}$ wavelength of maximal absorption; $\lambda_{em}$ wavelength of maximal emission; $\varepsilon$ molar absorptivity at $\lambda_{abs}$ (standard error is typically 10%); $\phi$ fluorescence quantum yield; molecular brightness = $\phi \times \varepsilon$; $K_D$ thermodynamic dissociation constant.

**Table S7.** Properties of the mutants isolated from the selections with HBP-3,5DM compared to FAST in PBS pH 7.4.

| Clone | Mutations (relative to FAST) | $\lambda_{abs}$ (nm) | $\lambda_{em}$ (nm) | $\varepsilon$ ($mM^{-1}cm^{-1}$) | $\phi$ | Molecular brightness | $K_D$ (µM) |
|---|---|---|---|---|---|---|---|
| FAST | | 464 | 522 | 30 | 0.16 | 4,000 | 3.7 |
| A-R5.3 | Q41H / V83L / M95I | 463 | 521 | 34 | 0.19 | 6,400 | 0.90 |
| A-R5.7 | Q41K / S72T / V83A / M95I | 463 | 525 | 27 | 0.23 | 6,100 | 0.53 |
| A-R5.12 | K17N / Q41K / M95T | 463 | 522 | 31 | 0.24 | 7,400 | 0.76 |
| A-R5.13 =shuffling mutant 4 | Q41L / S117I | 463 | 523 | 32 | 0.21 | 6,600 | 0.71 |
| A-R5.21 =shuffling mutant 5 | V83E / S117R | 463 | 522 | 33 | 0.20 | 6,400 | 0.69 |
| A-R6.2 | E93G / M95V / M109L | 464 | 525 | 30 | 0.20 | 5,800 | 0.94 |
| A-R7.1 | D20N / E81Q / V83I / M109L | 464 | 524 | 26 | 0.16 | 4,300 | 2.6 |
| A-R7.6 | G25R / A84S | 464 | 521 | 32 | 0.23 | 7,100 | 1.2 |
| A-R7.10 | Q32K / Y76N | 464 | 524 | 32 | 0.20 | 6,200 | 1.1 |
| A-R5.7-1 =shuffling mutant 1 | Q41K / S72T / V83A / M95I + M109L | 463 | 519 | 33 | 0.25 | 8,000 | 0.25 |
| A-R5.7-2 | Q41K / S72T / V83A / M95I + S117I | 463 | 521 | 36 | 0.26 | 9,000 | 0.36 |
| A-R5.7-3 | Q41K / S72T / V83A / M95I + M109L + S117I | 464 | 521 | 34 | 0.24 | 8,200 | 0.20 |
| A-R5.12-1 =shuffling mutant 2 | K17N / Q41K / M95T + S72T + V83A | 464 | 520 | 34 | 0.22 | 7,600 | 0.30 |
| A-R7.6-1 =shuffling mutant 3 | G25R / A84S + Q41K + M95T | 465 | 522 | 34 | 0.26 | 8,700 | 0.98 |
| B-R5.2 | K17N / Q41K / S72T / M95T | 463 | 521 | 30 | 0.24 | 7,000 | 0.35 |
| B-R5.11 | K17N / G25E / S72T / V83A / M95T | 463 | 521 | 32 | 0.27 | 8,700 | 0.31 |
| B-R5.19 | Q41K / K80M / A84S / N89D / M95T / M109L | 464 | 519 | 30 | 0.24 | 7,400 | 0.19 |
| B-R6.1 | K17N / G21E / G25R / A30V / Q41L / S72T / V83A / M95T / S117R | 464 | 520 | 36 | 0.27 | 9,800 | 0.17 |
| B-R6.10 | Q41K / S72T / V83A / S117R | 463 | 520 | 32 | 0.22 | 7,100 | 0.46 |
| B-R6.15 | N13S / Q41K / S72T / V83A / M95T | 464 | 519 | 29 | 0.24 | 7,000 | 0.32 |
| B-R6.19 | K17I / G25R / Q41K / A44T / K60R / V83A / N89D / E93K / M95T /M109L / S117R | 464 | 520 | 35 | 0.28 | 9,900 | 0.14 |
| B-R7.3 | K17N / Q41K / D65E / S72T / V83A / K106M / M109L | 464 | 520 | 35 | 0.23 | 8,000 | 0.30 |
| B-R6.1-1 = pFAST | K17N / G21E / G25R / A30V / Q41L / S72T / V83A / M95T / S117R + M109L | 465 | 520 | 37 | 0.27 | 10,000 | 0.15 |
| B-R6.1-2 | K17N / G21E / G25E / A30V / Q41L / S72T / V83A / M95T / S117R | 464 | 521 | 30 | 0.31 | 9,300 | 0.19 |
| B-R6.1-3 | K17N / G21E / G25R / A30V / Q4IK / S72T / V83A / M95T / S117R | 464 | 521 | 36 | 0.27 | 9,700 | 0.14 |

Abbreviations are as follows: $\lambda_{abs}$ wavelength of maximal absorption; $\lambda_{em}$ wavelength of maximal emission; $\varepsilon$ molar absorptivity at $\lambda_{abs}$ (standard error is typically 10%); $\phi$ fluorescence quantum yield; molecular brightness = $\phi \times \varepsilon$; $K_D$ thermodynamic dissociation constant.

**Table S8.** Properties of the mutants isolated from the selection with HBT-3,5DM compared to FAST in PBS pH 7.4.

| Clone | Mutations (relative to FAST) | $\lambda_{abs}$ (nm) | $\lambda_{em}$ (nm) | $\varepsilon$ ($mM^{-1}cm^{-1}$) | $\phi$ | Molecular brightness | $K_D$ (µM) |
|---|---|---|---|---|---|---|---|
| FAST | | 434 | 500 | 25 | 0.072 | 2,000 | 1.7 |
| shuffling mutant 1 | Q41K / S72T / V83A / M95I + M109L | 432 | 500 | 28 | 0.090 | 2,500 | 0.26 |
| shuffling mutant 2 | K17N / Q41K / M95T + S72T + V83A | 434 | 505 | 35 | 0.081 | 2,800 | 0.52 |
| shuffling mutant 3 | G25R / A84S + Q41K + M95T | 433 | 501 | 32 | 0.085 | 2,700 | 0.58 |
| shuffling mutant 4 | Q41L / S117I | 432 | 501 | 36 | 0.078 | 2,800 | 0.78 |
| shuffling mutant 5 | V83E / S117R | 433 | 500 | 31 | 0.075 | 2,300 | 0.62 |
| B-R6.11 | A16D / Q41K / Y76H / V83A / M95T / M109L / S117R | 435 | 497 | 27 | 0.10 | 2,700 | 0.23 |
| B-R6.13 = tFAST | G25R/ Q41K / S72T / A84S / M95A / M109L / S117R | 435 | 497 | 30 | 0.11 | 3,000 | 0.33 |
| B-R6.17 | A27V / Q41K/ V83A / M95T / M109L / S117I | 435 | 499 | 27 | 0.098 | 2,700 | 0.24 |
| B-R6.24 | K17N / G21E / G25R / A30V / Q41L / S72T / V83A / M95T / S117R | 434 | 497 | 29 | 0.094 | 2,700 | 0.26 |
| B-R7.2 | Q41K / K80R / V83A / M95T / M109L | 432 | 496 | 24 | 0.095 | 2,300 | 0.34 |
| B-R7.4 | Q41K / M95T / S117I | 432 | 496 | 26 | 0.095 | 2,500 | 0.53 |

Abbreviations are as follows: $\lambda_{abs}$ wavelength of maximal absorption; $\lambda_{em}$ wavelength of maximal emission; $\varepsilon$ molar absorptivity at $\lambda_{abs}$ (standard error is typically 10%); $\phi$ fluorescence quantum yield; molecular brightness = $\phi \times \varepsilon$; $K_D$ thermodynamic dissociation constant.

**Table S9.** Properties of the mutants isolated from the selection with HBP-3,5DOM compared to FAST in PBS pH 7.4.

| Clone | Mutations (relative to FAST) | $\lambda_{abs}$ (nm) | $\lambda_{em}$ (nm) | $\varepsilon$ ($mM^{-1}cm^{-1}$) | $\phi$ | Molecular brightness | $K_D$ (µM) |
|---|---|---|---|---|---|---|---|
| FAST | | 488 | 556 | 31 | 0.29 | 9,000 | 100 |
| shuffling mutant 1 | Q41K / S72T / V83A / M95I + M109L | 488 | 555 | 34 | 0.36 | 12,000 | 2.0 |
| shuffling mutant 2 | K17N / Q41K / M95T + S72T + V83A | 488 | 556 | 30 | 0.36 | 11,000 | 2.1 |
| shuffling mutant 3 | G25R / A84S + Q41K + M95T | 487 | 555 | 30 | 0.33 | 9,800 | 4.4 |
| shuffling mutant 4 | Q41L / S117I | 485 | 556 | 31 | 0.33 | 10,000 | 7.2 |
| shuffling mutant 5 | V83E / S117R | 488 | 556 | 32 | 0.30 | 9,500 | 5.9 |
| B-R6.7 | G7D / Q41K / S72T / A84S / M95I / M109L / S117R / R124L | 488 | 555 | 35 | 0.35 | 12,000 | 4.7 |
| B-R7.3 | K17N / Q41K / A84S / M95T / M109L / S117R | 487 | 555 | 33 | 0.37 | 13,000 | 2.8 |
| B-R7.7 | Q41K/ D48G / S72T / V83A / M95T / M109L / S117R | 485 | 555 | 37 | 0.30 | 11,000 | 2.7 |
| B-R7.22 | G21R / G25R / Q41K / Q60E / M95T / M109L / A112G | 490 | 557 | 33 | 0.34 | 12,000 | 5.9 |

Abbreviations are as follows: $\lambda_{abs}$ wavelength of maximal absorption; $\lambda_{em}$ wavelength of maximal emission; $\varepsilon$ molar absorptivity at $\lambda_{abs}$ (standard error is typically 10%); $\phi$ fluorescence quantum yield; molecular brightness = $\phi \times \varepsilon$; $K_D$ thermodynamic dissociation constant.

**Table S10.** Plasmids used in this study

| Plasmid | Expression host | Open reading frame | Ref. |
|---|---|---|---|
| pCTCON-FAST | Yeast | FAST | 2 |
| pAG681 | Yeast | shuffling mutant 1 (libraryA-R5.7-1 [HBP-3,5DM]) | this study |
| pAG682 | Yeast | shuffling mutant 2 (libraryA-R5.12-1 [HBP-3,5DM]) | this study |
| pAG683 | Yeast | shuffling mutant 3 (libraryA-R7.6-1 [HBP-3,5DM]) | this study |
| pAG684 | Yeast | shuffling mutant 4 (libraryA-R5.13 [HBP-3,5DM]) | this study |
| pAG685 | Yeast | shuffling mutant 5 (libraryA-R5.21 [HBP-3,5DM]) | this study |
| pAG686 | Yeast | pFAST | this study |
| pAG687 | Yeast | tFAST | this study |
| pAG688 | Yeast | oFAST | this study |
| pAG104 | Mammalian | FAST | 2 |
| pAG29 | Mammalian | EGFP | 2 |
| pAG490 | Mammalian | CMV-FRB-NFAST-IRES-mTurquoise2 | 3 |
| pAG244 | Mammalian | iFAST | 4 |
| pAG654 | Mammalian | pFAST | this study |
| pAG655 | Mammalian | tFAST | this study |
| pAG656 | Mammalian | oFAST | this study |
| pAG657 | Mammalian | H2B-pFAST | this study |
| pAG660 | Mammalian | lyn11-pFAST | this study |
| pAG665 | Mammalian | MAP4-pFAST | this study |
| pAG668 | Mammalian | LifeAct-pFAST | this study |
| pAG671 | Mammalian | mito-pFAST | this study |
| pAG897 | Mammalian | pDisplay-FAST | this study |
| pAG876 | Mammalian | pDisplay-pFAST | this study |
| pAG719 | Mammalian | MAP4-mVenus | this study |
| pAG849 | Mammalian | MAP4-Halotag | this study |
| X-888 | Mammalian/bird (CAG promoter) | Mito-pFAST | this study |
| X-858 | Mammalian/bird (CAG promoter) | H2B-pFAST | this study |
| X-889 | Mammalian/bird (CAG promoter) | H2B-FAST | this study |
| X-892 | Mammalian/bird (CAG promoter) | H2B-mCerulean | this study |
| pCX-H2B-EGFP | Mammalian/bird (CAG promoter) | H2B-EGFP | 5 |
| X-893 | Mammalian/bird (CAG promoter) | H2B-EYFP | this study |
| pCX-H2B-mKO | Mammalian/bird (CAG promoter) | H2B-mKO | 6 |
| X-158 | Mammalian/bird (CAG promoter) | Mb-EGFP | 7 |
| X-159 | Mammalian/bird (CAG promoter) | Mb-mCherry | 8 |
| X-736 | Mammalian/bird (CAG promoter) | Mb-iRFP670 | 9 |
| pCX-PACT-mKO | Mammalian/bird (CAG promoter) | PACT-mKO | 6 |

**Table S11.** Clones isolated in this study (pET28 plasmids for *E. coli* expression)

| Plasmid | Chromophore used for selection | Library | Clone | Mutations (relative to FAST) |
|---|---|---|---|---|
| pAG569 | HBO-3M | A | R6.8 | D65V / E93D / M109L / S117R |
| pAG562 | HBO-3M | A | R6.11 | V83I / M109L |
| pAG564 | HBO-3M | A | R7.1 | M109L |
| pAG565 | HBO-3M | A | R7.2 | Q41L / M109L |
| pAG566 | HBO-3M | A | R7.10 | V83I / T103I |
| pAG567 | HBO-3M | A | R7.12 | K17R / A30V / E74K / E93V |
| pAG568 | HBO-3M | A | R7.14 | T50S / E93Q / M95I |
| pAG562 | HBO-3,5DM | A | R6.6 | V83I / M109L |
| pAG563 | HBO-3,5DM | A | R6.18 | H3Q / D48A / T103I / M109L |
| pAG558 | HBO-3,5DM | A | R7.1 | N13I / D48E / D71N / V83G / M95V |
| pAG559 | HBO-3,5DM | A | R7.2 | Q41L / M95I / M109L |
| pAG560 | HBO-3,5DM | A | R7.5 | K60R / V83I / K104R / S117R |
| pAG561 | HBO-3,5DM | A | R7.7 | D71N / M109L / S117R |
| pAG795 | HBO-3,5DM | A-rational design | R7.7-1 | D71N / M109L / S117R + Q41L |
| pAG796 | HBO-3,5DM | A-rational design | R7.7-2 | D71N / M109L / S117R + Q41L + D48E |
| pAG644 | HBO-3,5DM | A-rational design | R7.7-3 | D71N / M109L / S117R + Q41L + D65V |
| pAG645 | HBO-3,5DM | A-rational design | R7.7-4 | D71N / M109L / S117R + Q41L + V83I |
| pAG797 | HBO-3,5DM | A-rational design | R7.7-5 | D71N / M109L / S117R + Q41L + M95I |
| pAG771 | HBP-3,5DM | A | R5.3 | Q41H / V83L / M95I |
| pAG772 | HBP-3,5DM | A | R5.7 | Q41K / S72T / V83A / M95I |
| pAG773 | HBP-3,5DM | A | R5.12 | K17N / Q41K / M95T |
| pAG556 | HBP-3,5DM | A | R5.13 | Q41L / S117I |
| pAG557 | HBP-3,5DM | A | R5.21 | V83E / S117R |
| pAG774 | HBP-3,5DM | A | R6.2 | E93G / M95V / M109L |
| pAG775 | HBP-3,5DM | A | R7.1 | D20N / E81Q / V83I / M109L |
| pAG776 | HBP-3,5DM | A | R7.6 | G25R / A84S |
| pAG777 | HBP-3,5DM | A | R7.10 | Q32K / Y76N |
| pAG553 | HBP-3,5DM | A-rational design | R5.7-1 | Q41K / S72T / V83A / M95I + M109L |
| pAG778 | HBP-3,5DM | A-rational design | R5.7-2 | Q41K / S72T / V83A / M95I + S117I |
| pAG779 | HBP-3,5DM | A-rational design | R5.7-3 | Q41K / S72T / V83A / M95I + M109L + S117I |
| pAG554 | HBP-3,5DM | A-rational design | R5.12-1 | K17N / Q41K / M95T + S72T + V83A |
| pAG555 | HBP-3,5DM | A-rational design | R7.6-1 | G25R / A84S + Q41K + M95T |
| pAG780 | HBP-3,5DM | B | R5.2 | K17N / Q41K / S72T / M95T |
| pAG639 | HBP-3,5DM | B | R5.11 | K17N / G25E / S72T / V83A / M95T |
| pAG781 | HBP-3,5DM | B | R5.19 | Q41K / K80M / A84S / N89D / M95T / M109L |
| pAG640 | HBP-3,5DM | B | R6.1 | K17N / G21E / G25R / A30V / Q41L / S72T / V83A / M95T / S117R |
| pAG782 | HBP-3,5DM | B | R6.10 | Q41K / S72T / V83A / S117R |
| pAG783 | HBP-3,5DM | B | R6.15 | N13S / Q41K / S72T / V83A / M95T |
| pAG642 | HBP-3,5DM | B | R6.19 | K17I / G25R / Q41K / A44T / K60R / V83A / N89D / E93K / M95T /M109L / S117R |
| pAG784 | HBP-3,5DM | B | R7.3 | K17N / Q41K / D65E / S72T / V83A / K106M / M109L |
| pAG641 | HBP-3,5DM | B-rational design | R6.1-1 | K17N / G21E / G25R / A30V / Q41L / S72T / V83A / M95T / S117R + M109L |
| pAG785 | HBP-3,5DM | B-rational design | R6.1-2 | K17N / G21E / G25E / A30V / Q41L / S72T / V83A / M95T / S117R |
| pAG786 | HBP-3,5DM | B-rational design | R6.1-3 | K17N / G21E / G25R / A30V / Q4IK / S72T / V83A / M95T / S117R |
| pAG791 | HBT-3,5DM | B | R6.11 | A16D / Q41K / Y76H / V83A / M95T / M109L / S117R |
| pAG643 | HBT-3,5DM | B | R6.13 | G25R/ Q41K / S72T / A84S / M95A / M109L / S117R |
| pAG792 | HBT-3,5DM | B | R6.17 | A27V / Q41K/ V83A / M95T / M109L / S117I |
| pAG640 | HBT-3,5DM | B | R6.24 | K17N / G21E / G25R / A30V / Q41L / S72T / V83A / M95T / S117R |
| pAG793 | HBT-3,5DM | B | R7.2 | Q41K / K80R / V83A / M95T / M109L |
| pAG794 | HBT-3,5DM | B | R7.4 | Q41K / M95T / S117I |
| pAG787 | HBP-3,5DOM | B | R6.7 | G7D / Q41K / S72T / A84S / M95I / M109L / S117R / R124L |
| pAG788 | HBP-3,5DOM | B | R7.3 | K17N / Q41K / A84S / M95T / M109L / S117R |
| pAG789 | HBP-3,5DOM | B | R7.7 | Q41K/ D48G / S72T / V83A / M95T / M109L / S117R |
| pAG790 | HBP-3,5DOM | B | R7.22 | G21R / G25R / Q41K / Q60E / M95T / M109L / A112G |

**Table S12.** Sequences of FAST and pFAST

| |
|---|
| **FAST (125 amino acids, MW = 13,706 Da)**<br>MEHVAFGSEDIENTLAKMDDGQLDGLAFGAIQLDGDGNILQYNAAEGDITGRDPKQVIGKNFFKDVAPGTDSPEFYGKFKEGVA<br>SGNLNTMFEWMIPTSRGPTKVKVHMKKALSGDSYWVFVKRV |
| **DNA sequence coding for FAST (375 bp)**<br>atggagcatgttgcctttggcagtgaggacatcgagaacactctggccaaaatggacgacggacaactggatgggttggcctttggcgcaattcagctcgatggtgacgggaatatc<br>ctgcagtacaatgctgctgaaggagacatcacaggcagagatcccaaacaggtgattgggaagaacttcttcaaggatgttgcacctggaacggattctcccgagttttacggcaa<br>attcaaggaaggcgtagcgtcagggaatctgaacaccatgttcgaatggatgataccgacaagcaggggaccaaccaaggtcaaggtgcacatgaagaaagccctttccggtg<br>acagctattgggtctttgtgaaacgggtg |
| **pFAST (125 amino acids, MW = 13,884 Da)**<br>MEHVAFGSEDIENTLANMDDEQLDRLAFGVIQLDGDGNILLYNAAEGDITGRDPKQVIGKNFFKDVAPGTDTPEFYGKFKEGAAS<br>GNLNTMFEWTIPTSRGPTKVKVHLKKALSGDRYWVFVKRV |
| **DNA sequence coding for pFAST (375 bp)**<br>atggagcatgttgcctttggcagtgaggacatcgagaacactctggccaatatggacgacgaacaactggataggttggcctttggcgtaattcagctcgatggtgacgggaatatcc<br>tgctgtacaatgctgctgaaggggacatcactggcagagatcccaaacaggtgattgggaagaacttcttcaaggatgttgcacctggaacggatactcccgagttttacggcaaatt<br>caaggaaggcgcagcgtcagggaatctgaacaccatgttcgaatggacgataccgacaagcaggggaccaaccaaggtcaaggtgcacttgaagaaagccctttccggtgac<br>agatattgggtctttgtgaaacgggtg |

**Table S13.** Imaging settings used in this study

| Figure | Panel | Fluorescent reporter | Excitation settings (nm) | Emission settings (nm) | Fluorescence microscopy | Comments |
|---|---|---|---|---|---|---|
| Fig. 2 | Panel e | oFAST:HBO-3,5DM | 405 | 450 - 550 | Confocal | |
| | Panel f | tFAST:HBT-3,5DM | 405 | 450 - 550 | | |
| | Panel g | pFAST:HBP-3,5DM | 488 | 500 - 600 | | |
| Fig. 4 | Panel a | pFAST:HBO-3M | 405 | 420 - 550 | Confocal | |
| | | pFAST:HBO-3,5DM | 405 | 450 - 550 | | |
| | | pFAST:HBT-3,5DM | 405 | 450 - 600 | | |
| | | pFAST:HBP-3M | 405 | 450 - 600 | | |
| | | pFAST:HBT-3,5DOM | 488 | 500 - 600 | | |
| | | pFAST:HBP-3,5DM | 488 | 493 - 600 | | |
| | | pFAST:HBP-3,5DOM | 488 | 495 - 680 | | |
| | | pFAST:HMBR | 488 | 495 - 600 | | |
| | | pFAST:HBRAA-3E | 488 | 490 - 700 | | |
| | | pFAST:HBRAA-3M | 488 | 490 - 700 | | |
| | | pFAST:HBRAA-3,5DM | 514 | 550 - 700 | | |
| | | pFAST:HBR-3,5DM | 488 | 500 - 630 | | |
| | | pFAST:HBR-3,5DOM | 514 | 550 - 700 | | |
| | | pFAST:HBIR-3,5DOM | 561 | 570 - 700 | | |
| | Panel b | pFAST:HBP3,5DM | 488 | 493 - 600 | Confocal | |
| | Panel c | pFAST:HBR-3,5DOM | 520 | 550 - 650 | | 81 optical sections ( z step 164 nm) |
| | Panel d | pFAST:HBR-3,5DOM | 520 | 550 - 650 | Confocal | Maximum intensity projection of 5 optical sections (for lyn11-pfast) |
| Fig. 5 | Panel b | pFAST:HBR-3,5DOM<br>mKO | 561 | 580 - 654 | Spinning-disk confocal | |
| | Panel c | pFAST:HBR-3,5DM<br>EYFP | 510/25 | 540/30 | Widefield | LED light source |
| | Panel d | pFAST:HBP-3,5DOM<br>EYFP | 470/24 | 540/30 | Widefield | LED light source |
| | Panel e | pFAST:HMBR<br>EGFP | 470/24 | 525/50 | Widefield | LED light source |
| | Panel f | pFAST:HBP-3,5DOM<br>EGFP | 470/24 | 525/50 | Widefield | LED light source |
| | Panel g | pFAST:HBP-3M<br>mCerulean | 440/20 | 480/40 | Widefield | LED light source |
| | Panel h | pFAST:HBR-3,5DOM | 561 | 580 - 654 | Spinning-disk confocal | |
| | | iRFP670 | 642 | 665 - 705 | | |
| | Panel i | pFAST:HBP-3,5DOM | 488 | 500 - 550 | | |
| | | mKO | 561 | 580 - 654 | | |
| | | pFAST:HMBR | 488 | 500 - 550 | | |
| | Panel j | mKO | 561 | 580 - 654 | | |
| | | iRFP670 | 642 | 665 - 705 | | |
| Fig. 6 | Panel a-l | pFAST:HBR-3,5DOM | 520 | 550 - 650 | 3D STED | 775 nm pulsed depletion laser ; motorized collar 93× glycerol NA 1.3 objective; optimized pixel size(from 25 to 45 nm) and an average line acquisition of 16 |
| Fig. S8 | Panel a,d | pFAST:HMBR | 488 | 495 - 600 | Confocal | |
| | Panel b | pFAST:HBR-3,5DOM | 514 | 550 - 700 | | |
| | Panel c | pFAST:HBR-3,5DM | 488 | 500 - 630 | | |
| | Panel e | pFAST:HBP-3,5DOM | 488 | 495 - 680 | | |
| | Panel f | pFAST:HBP-3,5DM | 488 | 493 - 600 | | |
| | Panel g | pFAST:HBP-3M | 458 | 470 - 580 | | |
| | Panel h | pFAST:HBT-3,5DOM | 488 | 495 - 600 | | |
| | Panel i | pFAST:HBT-3,5DM | 458 | 495 - 600 | | |
| Fig. S9 | Panel a | pFAST:HBRAA-3E | 488 | 490 - 700 | Confocal | |
| | | pFAST:HMBR | 488 | 490 - 700 | | |
| | Panel b | pFAST:HBRAA-3M | 488 | 490 - 700 | | |
| | | pFAST:HMBR | 488 | 490 - 700 | | |
| | Panel c | pFAST:HBRAA-3,5DM | 514 | 530 - 700 | | |
| | | pFAST:HMBR | 514 | 530 - 700 | | |
| Fig. S10 | Panel a | pFAST:HBR-3,5DOM<br>iFAST:HBR-3,5DOM | 488 | 550 - 700 | Confocal | |

| | | | | | | |
|---|---|---|---|---|---|---|
| | | FAST:HBR-3,5DOM | | | | |
| | | EGFP | | | | |
| | Panel b | pFAST:HBR-3,5DM | 488 | 500 - 650 | Confocal | |
| | | iFAST:HBR-3,5DM | | | | |
| | | FAST:HBR-3,5DM | | | | |
| | | EGFP | | | | |
| | Panel c | pFAST:HMBR | 488 | 495 - 600 | Confocal | |
| | | iFAST:HMBR | | | | |
| | | FAST:HMBR | | | | |
| | | EGFP | | | | |
| | Panel d,f,g | pFAST:HBP-3,5DOM | 488 | 495 - 600 | Confocal | |
| | | pFAST:HBP-3,5DM | | | | |
| | | EGFP | | | | |
| | Panel e,h,i | pFAST:HBT-3,5DOM | 458 | 470 - 600 | Confocal | |
| | | pFAST:HBP-3M | | | | |
| | | mTurquoise2 | | | | |
| Fig. S11 | | pFAST:HBP-3,5DM | 488 | 493 - 600 | Confocal | |
| Fig. S12 | Panel a | Halotag:TMR | 561 | 568 - 700 | Confocal | |
| | Panel b | mVenus | 514 | 520 - 600 | | |
| | Panel c | pFAST:HBP-3,5DM | 488 | 493 - 600 | | |
| Fig. S13 | | pFAST:HBR-3,5DOM | 520 | 550 - 680 | | |
| Fig. S15 | Panel a | pFAST:HBR-3,5DOM | 561 | 580 - 654 | Spinning-disk confocal | |
| | Panel b | pFAST:HBR-3,5DM | 510/25 | 540/30 | Widefield | LED light source |
| | Panel c | pFAST:HBP-3,5DOM | 470/24 | 540/30 | Widefield | LED light source |
| | Panel d | pFAST:HMBR | 470/24 | 525/50 | Widefield | LED light source |
| | Panel e | pFAST:HBP-3,5DM | 470/24 | 525/50 | Widefield | LED light source |
| Fig. S17 | Panel a | pFAST:HBR-3,5DM | 488 | 500 - 630 | Confocal | |
| | Panel b | pFAST:HBP-3,5DOM | 488 | 495 - 680 | | |
| | Panel c | pFAST:HMBR | 488 | 495 - 600 | | |
| | Panel d | pFAST:HBP-3,5DM | 488 | 495 - 680 | | |
| Fig. S18 | Panel a | pFAST:HBP-3,5DOM | 488 | 500 - 550 | Spinning-disk confocal | |
| | Panel b | pFAST:HMBR | 488 | 500 - 550 | | |
| Fig. S19 | Panel a,d | pFAST:HMBR | 488 | 495 - 550 | Confocal | Equipped with a spectral detector GaAsp of 32 channels (Airyscan module) |
| | Panel b | pFAST:HBR-3,5DOM | 561 | 570 - 750 | | |
| | Panel c | pFAST:HBR-3,5DM | 488 | 525 - 620 | | |
| | Panel e | pFAST:HBP-3,5DOM | 488 | 525 - 620 | | |
| | Panel f | pFAST:HBP-3,5DM | 488 | 495 - 650 | | |
| Movie S1 | | pFAST:HBR-3,5DOM | 561 | 580 - 654 | Spinning-disk confocal | |
| | | FAST:HBR-3,5DOM | | | | |
| | | EGFP | 488 | 500 - 550 | | |
| Movie S2 | | pFAST:HBR-3,5DOM | 561 | 580 - 654 | Spinning-disk confocal | |
| | | iRFP670 | 642 | 665 - 705 | | |
| Movie S3 | | pFAST:HBP-3,5DOM | 488 | 500 - 550 | Spinning-disk confocal | |
| | | mKO | 561 | 580 - 654 | | |
| Movie S4 | | pFAST:HMBR | 488 | 500 - 550 | Spinning-disk confocal | |
| | | mKO | 561 | 580 - 654 | | |
| | | iRFP670 | 642 | 665 - 705 | | |
| Movie S5 | | pFAST:HBR-3,5DM | 488 | 500 - 630 | Confocal | |
| | | pFAST:HBP-3,5DOM | 488 | 495 - 680 | | |
| | | pFAST:HMBR | 488 | 495 - 600 | | |
| | | pFAST:HBP-3,5DM | 488 | 495 - 680 | | |
| Movie S6 | | pFAST:HMBR | 488 | 500 - 550 | Spinning-disk confocal | |
| Movie S7 | | pFAST:HBR-3,5DOM | 561 | 570 - 750 | Confocal | Equipped with a spectral detector GaAsp of 32 channels (Airyscan module) Lyn11 : 1 image / 932 ms MAP4 : 1 image /4.99 s |
